# Poison exon splicing in the human brain: a new paradigm for understanding and targeting neurological disorders

**DOI:** 10.1101/2025.06.26.661736

**Authors:** Paolo Pigini, Huilin Xu, Yan Ji, Hannah Lindmeier, Henry R. Saltzman, Henry Shull, Shuqi Yun, Christiano R. R. Alves, Melissa A. Walker, M. Catarina Silva, Dadi Gao, Elisabetta Morini

## Abstract

**Background:** Poison exons (PE) are highly conserved exons whose inclusion creates premature termination codons (PTCs) and triggers nonsense-mediated decay (NMD) of the mature transcript. Despite their important role in post-transcriptional regulation, PEs remain poorly annotated due to the lack of systematic transcriptome-wide approaches.

**Results:** A comparative analysis of 957 eukaryotic transcriptomes revealed that *Homo Sapiens* exhibits the highest enrichment of NMD-targeted isoforms. By systematically predicting and annotating splicing events that generate PTCs, we found 9,814 PEs in the human genome. Using RNA-seq dataset from GTEx and BrainSpan, we analyzed PE splicing across tissues and developmental stages and identified 83 PEs uniquely found in the human brain, 973 PEs with brain-specific differential splicing, and 1,284 PEs differentially spliced during brain development. By integrating ClinVar variant annotations with SpliceAI predictions, we identified 1,625 pathogenic variants associated with neurological disorders that are predicted to affect the splicing of 743 PEs, and we functionally validated the impact of a subset of them using CRISPR prime-editing in human cells. Functional characterization of the deep intronic *NDUFAF6* c.420+784C>T variant associated with Leigh syndrome demonstrated increased PE inclusion and a significant reduction in NDUFAF6 protein levels in iPSC-derived induced neurons. Mutant cells also exhibited impaired mitochondrial membrane potential compared to controls. Together, these findings provide functional evidence that the c.420+784C>T variant promotes aberrant PE inclusion, leading to reduced NDUFAF6 expression and mitochondrial dysfunction.

**Conclusions:** Our findings highlight PEs as pivotal regulators of gene expression in the human brain and support the therapeutic targeting of PE splicing in neurological diseases.

## Background

Alternative mRNA splicing is a fundamental post-transcriptional regulatory mechanism that significantly expands the coding potential of eukaryotic genomes. Recent studies suggest that disruptions in this process contribute to approximately 10-30% of human genetic disorders (1–4), with nearly half of these alterations occurring in the intronic regions (5). Alternative splicing is particularly important in the brain, where it plays critical roles in neurodevelopment, neuronal differentiation, and synaptic function (6–11). This regulatory mechanism helps define the differentiation trajectories of various cell types within the nervous system (9, 12, 13) and appears to have undergone substantial evolutionary diversification in vertebrates, especially primates (14). Consequently, splicing alterations have been increasingly implicated in the pathogenesis of neurological disorders (15, 16), with non-coding variants emerging as particularly significant contributors (17).

Poison exons (PE) are naturally occurring, evolutionarily conserved exons that introduce premature termination codons (PTCs) when included in the mature transcript, thereby triggering nonsense-mediated decay (NMD) of the mature transcript and reducing protein expression (18–25). PEs act as finely tuned post-transcriptional regulators of gene expression (26) and have been implicated in both physiological and pathological processes (27). In murine models, PE-induced NMD has been proposed to downregulate non-neuronal genes during brain development (28). In humans, they contribute to key cellular functions, including lineage specification, neuronal differentiation, and neuroprotection (7, 29), and are also linked to disease mechanisms such as cancer (27). In the human brain, PEs help establish cellular identity during differentiation and play critical roles in the temporal and cell type–specific regulation of synaptic proteins, neuron-specific RNA-binding proteins (24, 30, 31), and chromatin modifiers (32). Notably, the splicing of PEs in the brain is regulated by a network of RNA-binding splicing regulators, including *RBFOX1*, *PTB1* and *NOVA1* (6, 7, 31, 33).

The dysregulation of PE splicing has been implicated in various neurological disorders (28). For example, aberrant inclusion of the PE “20N” in the *SCN1A* gene is associated with Dravet syndrome (DS) (34–37). An intronic variant disrupting normal skipping of a PE in the *FLNA* gene was found in a family segregating with periventricular nodular heterotopia (PVNH), a condition caused by abnormal neuronal migration during fetal brain development (7). In amyotrophic lateral sclerosis (ALS), *TDP-43* gene dysfunction disrupts PE regulation in iPSC-derived neurons, potentially contributing to disease pathogenesis (38, 39). Beyond their emerging role in disease etiology, PEs are also increasingly recognized as potential novel therapeutic targets for the treatment of neurological disorders (40–43).

Recent advances in computational approaches have improved identification of both PEs (44) and pathogenic variants affecting their splicing (2, 45). However, the field still lacks a comprehensive catalog of PE splicing events (46–48). In this study, we first conducted an evolutionary analysis by predicting NMD-targeted transcripts based on genome annotations, and found that *Homo sapiens* exhibits a striking enrichment of genes expressing transcripts containing PEs. We leveraged large RNA-seq datasets from GTEx and BrainSpan projects to identify human brain-specific and neurodevelopmentally regulated PEs. By integrating ClinVar variant annotations with SpliceAI predictions, we identified pathogenic variants that disrupt PE splicing and are associated with neurodevelopmental and neurodegenerative disorders. To functionally validate the impact of a subset of these variants on PE splicing, we applied an optimized CRISPR prime editing workflow (49) in human cell lines. Together, this work provides new insights into the role of PEs in brain function and disease, and highlights their potential as therapeutic targets for neurological conditions.

## Results

### The human transcriptome exhibits a unique enrichment of predicted NMD isoforms

Conventionally, NMD mRNA isoforms are defined by the presence of a PTC located more than 50 base pairs (bp) upstream of the last exon–exon junction (21–24) (Fig. 1A). Using these criteria, we systematically identified NMD isoforms across a dataset of 957 annotated eukaryotic genomes from Ensembl (Release 60 and 113), spanning the major clades of vertebrates, invertebrates, fungi, protists, and plants (Fig. 1B and Supplementary Table S1). This large-scale dataset enabled us to evaluate the evolutionary distribution and potential functional relevance of NMD across eukaryotes. On average, 2.67% of coding genes per genome were predicted to produce at least one NMD isoform (or NMD-regulated genes), though this proportion varied substantially across evolutionary lineages. Vertebrates (3.7%) and plants (3.7%) exhibited the highest fraction of coding genes producing at least one NMD isoform, followed by invertebrates (1.9%), while lower percentages were observed in fungi (0.6%) and protists (0.1%) (Fig. 1B and Supplementary Fig. S1A). A similar trend was observed when analysing the percentage of NMD isoforms over all isoforms in each species, with vertebrates (2.4%), invertebrates (2.0%), and plants (3.7%) showing higher values compared to protists (0.1%) and fungi (0.7%) (Supplementary Fig. S1B). We further evaluated the average percentage of NMD isoforms per gene in each species. In this case, we found that vertebrates (60.5%) and invertebrates (80.2%) display the highest values, followed by plants (45.6%), fungi (30.3%) and finally protists (18.1%) (Supplementary Fig. S1C). These data suggest that alternative splicing leading to NMD is particularly prominent in multicellular lineages with complex developmental processes. Among all species analysed, *Homo sapiens* emerged as the species with the highest number of predicted NMD isoforms. We found that 39.6% of human protein-coding genes give rise to at least one predicted NMD isoform.

**Figure 1.**
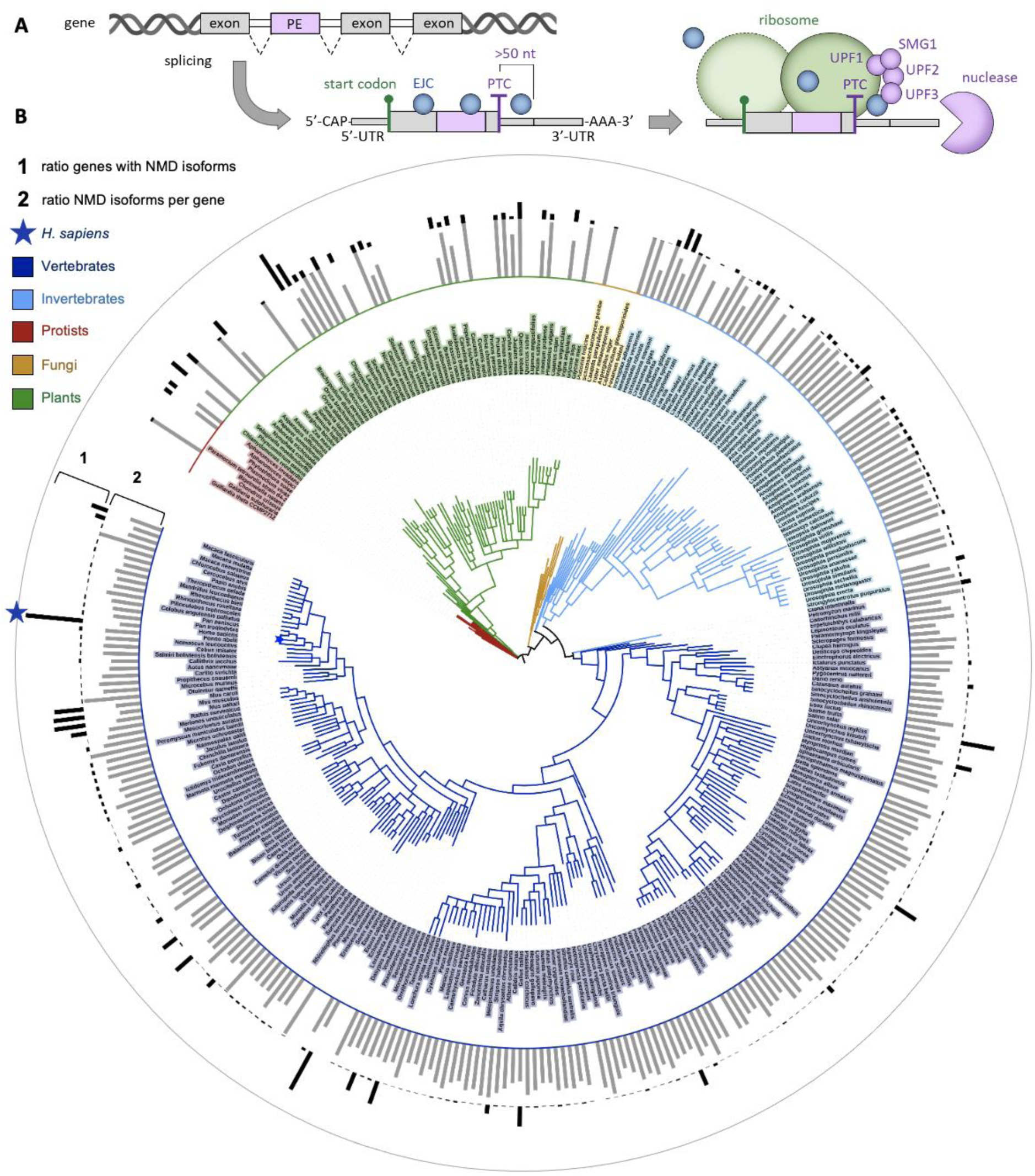
Evolutionary analysis of NMD isoforms across eukaryotes. (**A**) Schematic representation of a transcript carrying a PTC located more than 50 nt upstream of the last exon-exon junction, which triggers NMD. The ribosome stalls at the PTC and, unable to displace the exon-junction complex (EJC), triggers NMD via factors UPF1, UPF2, UPF3 and SMG1. (**B**) NMD isoforms annotated throughout the genomes of 957 eukaryotic species among vertebrates, invertebrates, protists, fungi and plants. For each species, bars indicate either the ratio of coding genes with at least one annotated NMD isoform (Track “1”, highest value: 0.40), or the average ratio of annotated NMD isoforms over all isoforms of each coding gene (Track “2”, highest value: 1.00).

Furthermore, in *Homo sapiens*, NMD isoforms represent 20.1% of all protein-coding gene isoforms, while an average of 35.0% of isoforms for a given protein-coding gene are predicted to undergo NMD. To account for differences in annotation depth between *Homo sapiens* and other species, we repeated the analysis using more stringent criteria. First, we restricted the species to those with high annotation completeness (completeness score ≥ 90%), as assessed by BUSCO (50). Second, we filtered the *Homo sapiens* annotation to include only transcripts with a “transcript support level” (TSL) of 1-3, thereby retaining only isoforms with strong experimental support as defined by the Ensemble project. Even under these more stringent conditions, *Homo sapiens* remains among the species with the highest proportion of predicted NMD isoforms. Specifically, 30.3% of human protein-coding genes contain at least one predicted NMD isoform, 14.3% of all isoforms from protein-coding genes are predicted to undergo NMD, and, on average, 38.6% of isoforms per protein-coding gene are classified as NMD-sensitive. Together, these results indicate that these observations are not driven by uneven annotation depth or the inclusion of low-confidence transcripts (Supplementary Fig. S1D and Supplementary Table S1 for the full dataset). Instead, they suggest that NMD plays a prominent role in regulating transcript stability in *Homo sapiens*, potentially reflecting the complexity and regulatory precision of human gene expression.

To explore the evolutionary trend of NMD occurrence, we investigated whether it could be associated with the complexity of the genome and transcriptome. Surprisingly, we observed no striking correlation between the total number of coding genes and the percentage of NMD-regulated genes across species (Supplementary Fig. S2A) or the frequency of NMD isoforms (Supplementary Fig. S2B and S2C). Notably, vertebrates often displayed higher percentages of NMD-regulated genes despite having fewer genes overall compared to plants, suggesting that genome size alone does not explain NMD frequency. In contrast, we found a positive association between transcriptome complexity and NMD abundance. Species with a greater number of isoforms per gene tend to have more NMD-regulated genes (p < 0.0001, Supplementary Fig. S2D) and a higher overall abundance of NMD isoforms (p < 0.0001, Supplementary Fig. S2E). However, this increase is driven primarily by a larger number of genes containing at least one NMD isoform, rather than by an increase in the average number of NMD isoforms per gene (Supplementary Fig. S2F). This pattern may reflect the fact that higher occurrence of alternative splicing may naturally increase the probability of producing NMD isoforms. We next examined whether exon density correlates with NMD occurrence. We observed a positive correlation (p ≤ 0.01 in all comparisons) between the average number of exons per gene and the number of NMD-regulated genes (Supplementary Fig. S2G) and the frequency of NMD isoforms (Supplementary Fig. S2H and S2I), consistent with the idea that splicing complexity increases opportunities for NMD-generating splicing events. Finally, we found a significant positive correlation (p < 0.0001 in all comparisons) between the average transcript length in each species and both the ratio of NMD-regulated genes (Supplementary Fig. S2J) and the frequency of NMD isoforms (Supplementary Fig. S2K and S2L). This could be explained by the previously observed correlation between NMD occurrence and the number of exons per gene.

Our evolutionary analysis highlights that NMD isoforms are broadly expressed across eukaryotes and are particularly enriched in vertebrates, especially in *Homo sapiens*. *Homo sapiens* exhibits the highest density of NMD events among all surveyed species. Our findings also suggest that the occurrence of NMD isoforms has been favoured by increasingly complex transcriptomes over evolution.

### Comprehensive classification of alternative splicing events involving PEs in human transcriptome

Building on the finding that *Homo sapiens* exhibits a high frequency of NMD isoforms, we next sought to systematically annotate the full repertoire of PEs in the human transcriptome. PEs are highly conserved exon cassettes whose inclusion introduces PTCs in the mature transcript, thereby triggering NMD and reducing protein levels. To identify these exons at scale, we developed an *in silico* strategy using genomic sequences and exon annotations (Supplementary Fig. S3A, see Methods and Supplementary Methods). We defined a PE as an exon that: (1) is ≤ 300 bp in length; (2) does not overlap the 5’ or 3’ UTR region of its host gene; and (3) upon inclusion into an annotated protein-coding transcript, generates a PTC located more than 150 bp downstream of the transcription start site and more than 50 nucleotides upstream of the last exon–exon junction, while remaining outside of the terminal exon (see Supplementary Methods for details). This stringent definition of PTC location was specifically chosen to minimize the likelihood of NMD escape (51), a mechanism by which some NMD transcripts avoid degradation. Using this approach, we identified 9,814 putative PEs, accounting for 17.8% of 55,150 annotated alternative splicing events extracted from the Gencode v26 GTF annotation, each of which is predicted to generate an NMD-sensitive isoform upon inclusion (Fig. 2A and Supplementary Table S2). Importantly, predicted PEs represent context-dependent splicing events that introduce a bona fide PTC only in the context of the identified transcript and coding frame. A comparative analysis across model organisms, including *Mus musculus*, *Danio rerio*, *Drosophila melanogaster*, *Caenorhabditis elegans*, *Saccharomyces cerevisiae*, and *Arabidopsis thaliana*, revealed species-specific differences in PE frequency, with *Homo sapiens* having the highest number of putative PE splicing events (Fig. 2A and Supplementary Table S3).

**Figure 2.**
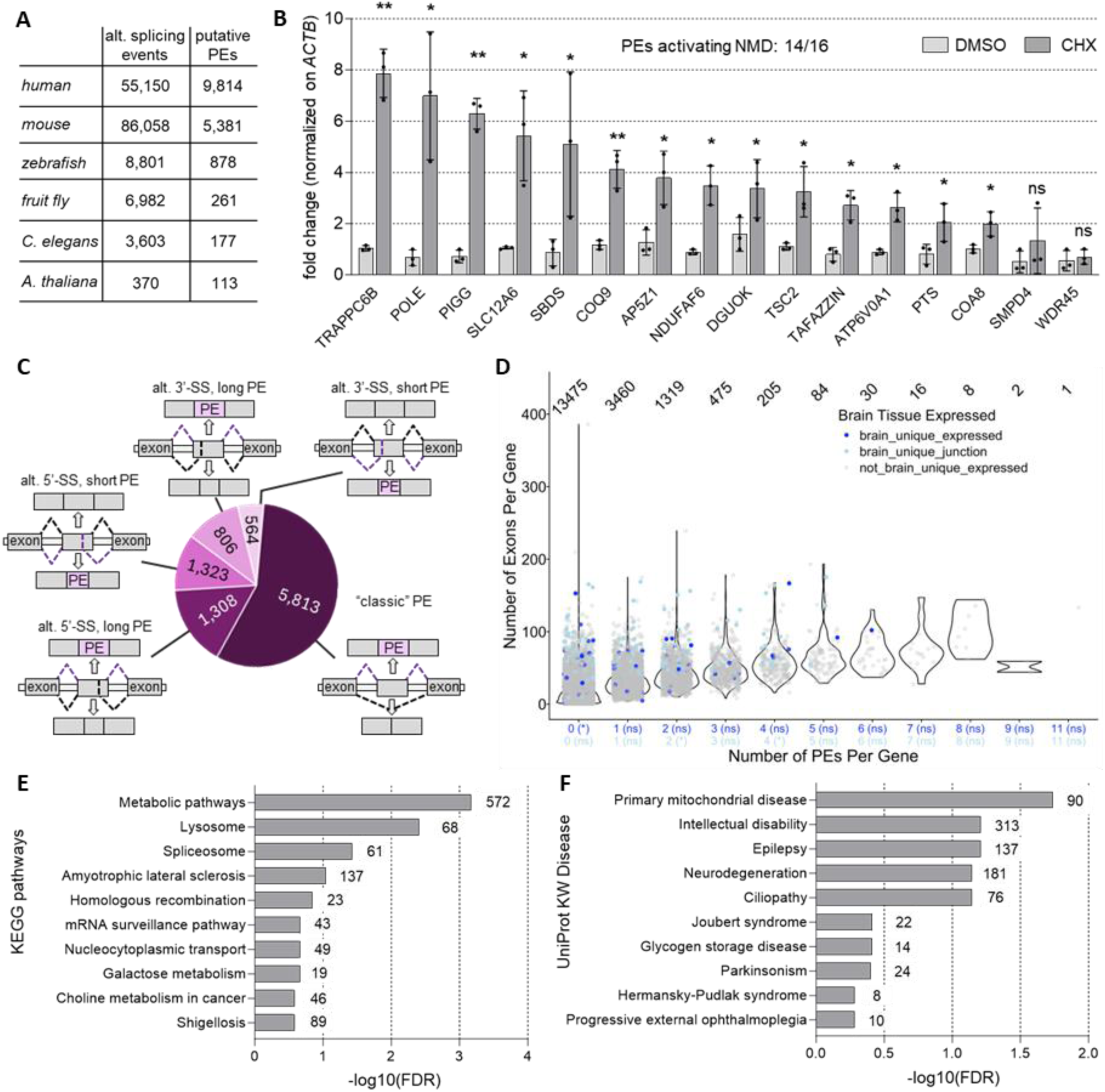
Analysis of putative PE. (**A**) Alternative splicing events involving putative PEs are counted for the genome annotation of different biological models. (**B**) Sensitivity to NMD assessed by CHX treatment in HEK293T cells for 16 putative PEs. Cells were treated with 50 µg/mL CHX (or DMSO), and the increase of transcript levels was evaluated by RT-qPCR. All values are internally normalized on *ACTB*, while transcript levels are represented as fold change compared to the DMSO. Statistical significance by Student’s t-test is shown for every comparison. * p<0.05, ** p<0.01, *** p<0.001, **** p<0.0001, ns: not significant. Data are shown as average ± standard deviation. Each data point represents one of three replicates. (**C**) Different types of alternative splicing events among putative PEs in *Homo sapiens*. They include canonical PE exclusion or inclusion (“classic”) events and alternative 3’- and 5’-splice-sites (“alt. 3’-SS” and “alt. 5’-SS” respectively). Labels “long” and “short” refer to whether the longer or the shorter isoform of the two contain the PE (see Supplementary Methods for details). (**D**) Number of unique putative PEs counted for each annotated gene in the human genome (x-axis) against the total number of exons in each gene (y-axis). Genes are divided into three groups (see Methods): 1) only expressed in the brain (dark blue); 2) expressed in multiple tissues but have a splice junction only used in the brain (light blue); and 3) others (grey). The brackets on the x-axis indicate whether PE- inclusion is significantly enriched in gene groups. (**E**) Pathway and (**F**) disease enrichment analysis by DAVID tool for human genes that include PEs. Numbers indicate the total count of genes identified in each category.

To evaluate the robustness of our PE identification, we then assessed *in vitro* whether their inclusion indeed triggers NMD by measuring transcript levels after treating human cells with cycloheximide (CHX), which inhibits translation and stabilizes NMD-targeted mRNAs (28, 52, 53). We performed this analysis in HEK293T cells for a subset of identified transcripts containing putative PEs. The 16 PEs selected for validation were chosen based on the computational predictions and expression detectability. Reverse-transcription quantitative PCR (RT-qPCR) following CHX or DMSO treatment revealed that 87.5% of transcripts containing PEs were significantly stabilized by CHX, confirming their degradation via NMD (Fig. 2B and Supplementary Fig. S3B). Transcripts without CHX-induced upregulation likely represent cases in which downstream splicing events restore the reading frame. Specifically, in *SMPD4* and *WDR45* the predicted PEs generate a PTC only in specific transcript isoforms that are not expressed in HEK293T cells. In these cells, inclusion of adjacent exons restores the open reading frame, thereby preventing NMD activation explaining the lack of CHX responsiveness (Supplementary Fig. S3C).

More than half of these PE splicing events (5,813 events, or 59.2%) arise from cassette exons, while the remaining result from alternative 3′ or 5′ splice site usage (Fig. 2C and Supplementary Fig. S3D). PEs did not show a preferential position along the gene (Supplementary Fig. S3E). While genes expressed in the brain, particularly those involved in neuronal and synaptic functions, are well-known to be longer and to contain more exons (54–56), we did not observe them to contain a higher number of PEs per gene (Fig. 2D). In fact, brain-expressed genes had fewer PEs than non-brain genes (Fig. 2D), suggesting a requirement for strict PE regulation in the brain transcriptome.

To gain insight into the biological functions of genes producing transcripts with PEs, we conducted pathway and disease enrichment analyses. KEGG pathway analysis revealed a significant enrichment (FDR = 0.00069, 572 genes) in metabolic pathways (Fig. 2E and Supplementary Table S4). We also found an enrichment for functions related to the spliceosome (61 genes) and mRNA surveillance (43 genes). Several splicing regulators were among the enriched genes, such as SRS (57) factors and SF3 (58) factors. These findings are consistent with the well-established data indicating that many splicing factors regulate their own expression through negative feedback loops mediated by PE inclusion (19, 27, 59). Additionally, disease ontology enrichment using the UniProt KW Disease database (Fig. 2F and Supplementary Table S5) identified a preferential enrichment of genes containing PE in neurological disorders, most prominently epilepsy (137 genes), intellectual disability (313 genes) and neurodegeneration (181 genes), reinforcing the hypothesis that PE splicing plays a key regulatory role in brain functions (24, 30, 31).

### PEs exhibit brain-specific and spatio-temporal regulation during neurodevelopment

To investigate the spatiotemporal regulation of PE splicing in the human brain, we leveraged large-scale transcriptomic datasets from both adult and developing tissues. We hypothesized that dynamic inclusion or exclusion of PEs may contribute to brain-specific gene regulation. We first analysed RNA-seq data from the Genotype-Tissue Expression (GTEx) Project, which encompasses 17,382 libraries derived from 54 distinct tissues, including 13 brain regions from 838 donors without major chronic diseases (60). This analysis (see Methods) revealed that 973 PEs are differentially spliced in the brain compared to other tissues (Fig. 3A, Supplementary Fig. S4, and Supplementary Table S6), and 83 PEs are uniquely detected in the brain. As anticipated, inclusion of these PEs correlated inversely with expression of their host genes (Supplementary Fig. S5), consistent with their role in post-transcriptional downregulation via NMD. To explore the functional implications of these brain-regulated PEs, we conducted gene ontology (GO) analyses by stratifying them based on PSI values. Using 0.5 as a PSI cut-off, we could divide genes into low PE inclusion (PSI < 0.5) and high PE inclusion (PSI > 0.5), and found that genes showing low PE inclusion in the brain are significantly enriched in RNA metabolism and splicing-related pathways (Fig. 3B and Supplementary Table S7). Importantly, expression of genes with high PE inclusion was still detectable, likely reflecting the well-documented incomplete efficiency of NMD (47, 61–69).

**Figure 3.**
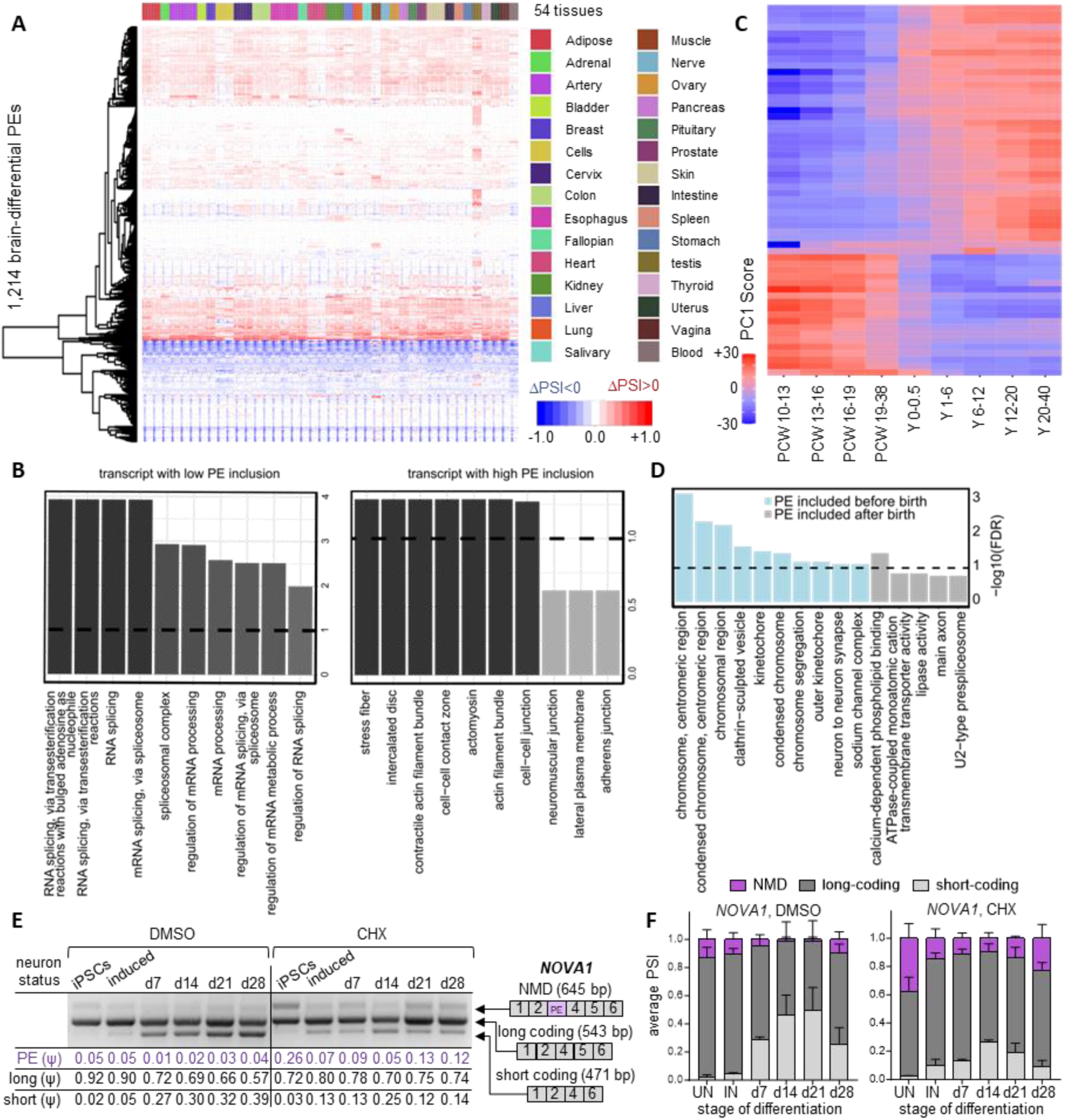
Analysis of PE splicing in the human brain. (**A**) Heatmap showing differential PSI (ΔPSI) of PEs across 54 GTEx tissues relative to brain. Each column represents a tissue, and each row represents one of the 973 PEs identified as differentially spliced in the brain. Negative values indicate higher PE inclusion in the brain, while positive values indicate lower inclusion. Hierarchical clustering is based on ΔPSI values. (**B**) GO and pathway enrichment analysis for host genes of with low PE inclusion (PSI < 0.5) or high PE inclusion (PSI > 0.5) in the brain. The top 10 enrichment groups (ranked by statistical q value) are shown for each analysis. (**C**) Heatmap of differential splicing of PEs across brain developmental stages (BrainSpan database). Each column represents a developmental stage, and each row represents a PE cluster (grouped by similarity of GAM-filtered trajectories, see Methods). Colours indicate splicing change (represented as PC1 trajectory value), with positive values indicating increased PE inclusion and negative values indicating decreased inclusion (see Methods). (**D**) GO and pathway enrichment analysis for host genes of PEs showing higher splicing inclusion before (light blue) or after (grey) birth. The top enrichment groups with FDR < 0.1 are shown for each analysis. (**E**) Splicing analysis of a brain-specific PE in the *NOVA1* gene across neuronal differentiation, including undifferentiated iPSCs, induced neurons following 3 days of NGN2 overexpression, and neurons at 7, 14, 21 or 28 days of maturation. Splicing isoforms corresponding to each band are indicated on the right, and band identity was confirmed by Sanger sequencing. Cells were treated either with DMSO (vehicle) or 50 μg/ml CHX. (F) PSI values of the different *NOVA1* isoforms were calculated by densitometry analysis of RT-PCR bands from three different replicates. Cells were treated either with DMSO (vehicle) or 50 μg/ml CHX.

We then analysed the developmental regulation of PEs using the BrainSpan Atlas, a comprehensive resource comprising 607 libraries from 41 individuals across 16 distinct brain regions. These libraries span key developmental stages, ranging from 10 post-conception weeks (pcw) to 40 years of age. This analysis (see Methods) identified 1,284 differentially spliced PEs in at least one of the 16 brain regions (Supplementary Fig. S6-8, Supplementary Table S8). Clustering of these PEs revealed distinct developmental splicing patterns, characterized by higher PE inclusion before or after birth (Fig. 3C). GO enrichment of the clustered PEs with higher inclusion before birth revealed significant associations with cell growth and division, while clustered PEs with higher inclusion after birth showed involvement in calcium-dependent phospholipid binding (Fig. 3D and Supplementary Table S9). It is worth noting that the PE inclusion showed predominantly non-linear trajectories across development and exhibited region-specific patterns (Supplementary Fig. S9A-P and Supplementary Table S10).

Finally, our analyses showed that splicing factors, including *RBFOX1*, *NOVA1*, and *UPF3B,* were significantly enriched for brain-specific PEs (p = 0.000004, one-sided hypergeometric test). We then investigated whether PE inclusion in the transcripts of these splicing factors indeed triggers NMD and changes during neuronal development, using iPSC-derived i3N neurons, a model of human neurons that differentiate via *NGN2* induction (70). We specifically selected this neuronal cell model because the PSI values of the PE splicing better mirror those observed in the human brain (Supplementary Fig. S10A). After *NGN2*-mediated induction and a differentiation period of up to 28 days (Supplementary Fig. S10B and S10C), we assessed PE splicing by RT-PCR. We found that PE inclusion in *NOVA1* was higher in iPSCs but declined during neuronal maturation, coinciding with increased expression of coding isoforms (71), which was more evident after CHX treatment (Fig. 3E and 3F). For splicing factor *TARDBP* we were able to validate that the inclusion of the PE triggered NMD, although no significant differentiation-specific splicing changes were observed, while for *RBFOX1* and *UPF3B* we could not detect the presence of the respective PEs at any differentiation stages (data not shown). Of note, consistently with the result in HEK293T cells, analysis of CHX-treated neurons confirmed that for the majority of putative PEs (87.5%) indeed trigger NMD (Supplementary Fig. S11).

Our data reveal that PE splicing is highly dynamic across brain regions and developmental stages, suggesting a widespread mechanism of post-transcriptional regulation in the nervous system. The enrichment of PE-regulated genes in RNA metabolism and neurodevelopmental pathways, along with validation of key neuronal splicing regulators, supports a model in which PEs act as molecular switches to fine-tune gene dosage with temporal and spatial precision.

### Identification of pathogenic variants associated with neurological disorders and disrupting PE splicing

To investigate the role of PE mis-splicing as a pathogenic mechanism for brain disorders, we sought to identify variants affecting PE splicing. We analysed all pathogenic and likely pathogenic variants in the ClinVar database (72) and used SpliceAI (2) to predict their impact on mRNA splicing. As expected, most pathogenic and likely pathogenic variants predicted to impact mRNA splicing (SpliceAI score > 0.2) were located in intronic regions: 45,691 intronic versus 22,991 exonic variants (Supplementary Fig. S12). For each variant, we assigned a positive or negative SpliceAI score to indicate the direction of the effect: positive scores indicate that the variant increased PE inclusion, and negative scores indicate that the variant decreased PE inclusion (significant absolute value > 0.2). We then compared these predictions with PSI values from healthy brain tissues (GTEx) to prioritize variants significantly affecting PE splicing in the brain. We specifically considered two scenarios (see Methods): (1) a PE with a PSI < 0.7 in the wild-type, for which the variant is predicted to increase inclusion; and (2) a PE with a PSI > 0.3 in the wild-type, for which the variant is predicted to decrease inclusion. Using this approach, we identified 1,625 pathogenic variants predicted to affect the splicing of 743 PEs (Fig. 4A and Supplementary Table S11). We then mapped these variants to associated diseases and clinical phenotypes using the ODiseA ontology. Notably, a large fraction of these variants was associated with seizures (596 variants) and epileptic syndromes (262 variants), including DS, consistent with previous findings of PE mis-splicing as a pathogenetic mechanism in these disorders^22,26^ (Fig. 4B). In addition, we observed enrichment in genes associated with various neurodevelopmental (e.g. Joubert syndrome and microcephaly) and neurodegenerative conditions (e.g. Alzheimer’s disease and Parkinson’s disease). These findings suggest that altered PE splicing may represent a common mechanism contributing to a wide range of neurological disorders. To further assess the robustness of these predictions, we performed a comparative analysis using Pangolin (45), a deep learning-based splicing predictor with an output structure comparable to that of SpliceAI. We re-analysed all pathogenic and likely pathogenic ClinVar variants using Pangolin, and observed strong concordance with SpliceAI for overlapping predictions (Supplementary Fig. S13), supporting the robustness of our approach. At the same time, SpliceAI showed higher sensitivity, identifying additional candidate events, while Pangolin uniquely identified a smaller subset. Together, these results support the reliability of our pipeline while highlighting the value of integrating multiple predictors.

**Figure 4.**
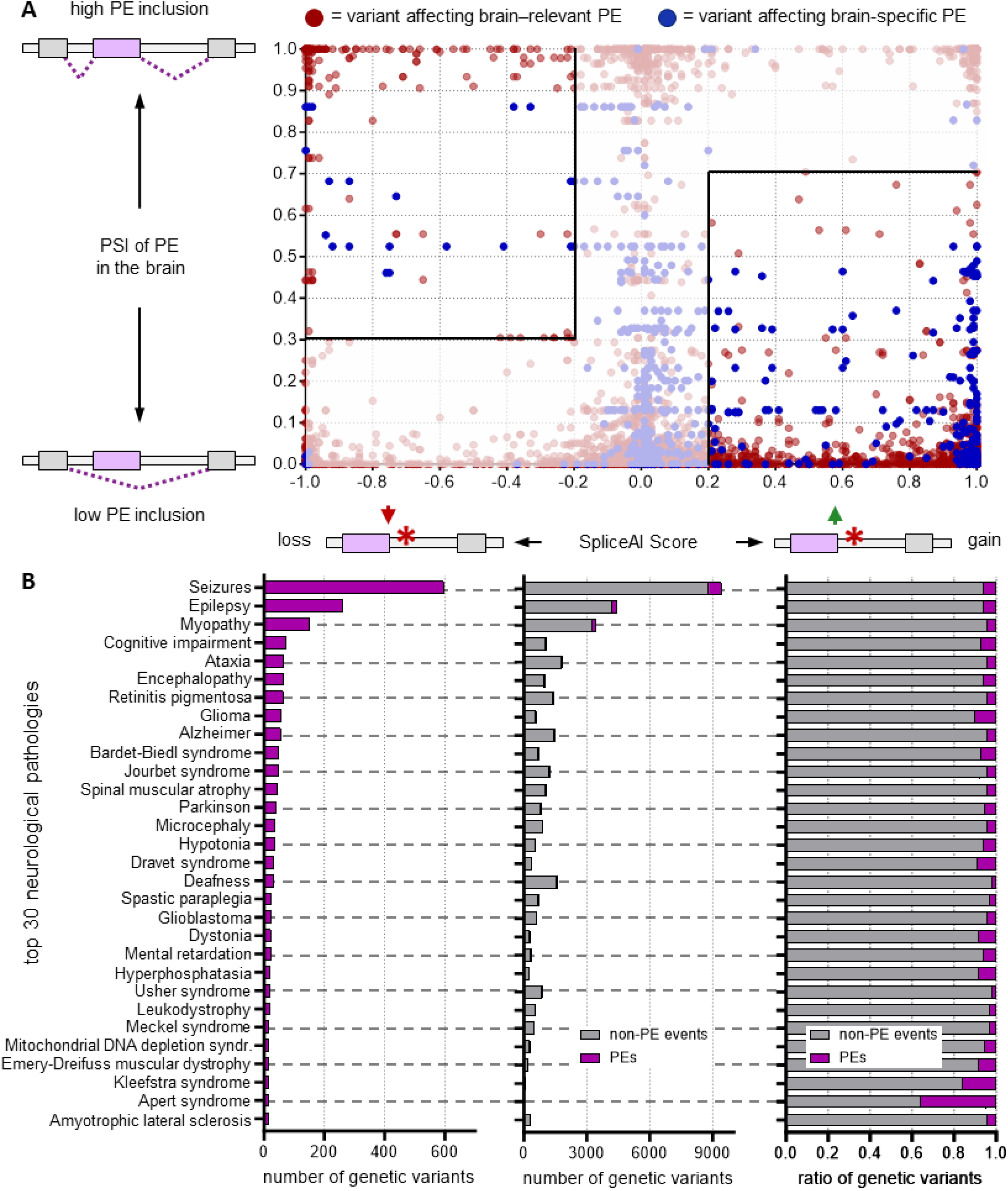
Pathogenic variants affecting PE splicing in neurological disorders. (**A**) Prioritizing pathogenic ClinVar variants based on its SpliceAI scores (x-axis) and the PSI values of PE splicing event in healthy human brain(y-axis). Each dot represents a variant/PE match. Blue dots represent PEs that are differentially spliced in the brain, while red dots represent PEs whose host gene is involved in neuronal functions as per GO. PSI were calculated in healthy brain tissues from the GTEx database. SpliceAI score have been assigned with a positive or a negative value depending on whether the mutation increases or decreases PE inclusion, respectively. Highlighted areas indicate variants that are highly likely to promote PE exclusion (GTEx PSI>0.7, SpliceAI score<-0.2) or inclusion (GTEx PSI<0.3, SpliceAI score>+0.2). (**B**) Neurological disorders associated with pathogenic variants affecting PE splicing are ranked by the number of such variants. Disease associations come from GO analysis and the ODiseA pathology database. The left panel shows the number of genetic variants affecting PEs. The other two panels demonstrate the number (middle) and the ratio (right) of variants affecting PEs, compared to the number of those affecting non-PE splicing events. In all panels, pathologies are ranked based on the number of variants affecting PE splicing.

To experimentally validate the effects of pathogenic variants on PE splicing, we developed and optimized a CRISPR prime editing workflow (Supplementary Fig. S14A, see Methods) (49, 73–75). As proof-of-concept, we designed and tested 13 pegRNAs to introduce variants predicted to affect PE splicing in 12 different genes. Editing efficiency was assessed in HEK293T cells, with 10 of the 13 achieving >5% efficiency, and 9 exceeding 10% (average: 22.9%; Supplementary Fig. S14B). The effect of selected variants on PE splicing was assessed by RT-PCR in cells treated with CHX, allowing visualization of PE splicing events that would otherwise be reduced by NMD. In parallel, we also performed RT-qPCR to quantitatively measure changes in PE inclusion levels, providing a robust assessment of both the direction and magnitude of the splicing alterations (Supplementary Fig. S14C). Importantly, in all selected genes, the introduction of the pathogenic variant produced the predicted change in PE inclusion (Supplementary Fig. S14C). Variable editing efficiencies across prime-edited cell populations may partially attenuate the observed magnitude of these effects; however, the consistent directionality of the changes across all tested variants supports the robustness of the predictions. Here, we report five representative examples from the selected variants (Fig. 5 and Supplementary Fig. S15), selected based on clinical relevance. Variants in these genes were predicted to have a distinct effect on PE splicing. Two mutations were predicted to increase PE inclusion. Of these, *TAFFAZIN* (or *TAZ*) c.284+110G>A, located in intron 3 (76), was predicted to enhance PE inclusion(SpliceAI score: 0.57) and result in a +0.12 PSI change in a bulk-edited population with an average of 15% editing efficiency (Fig. 5A). *NDUFAF6* c.420+784C>T, associated with Leigh syndrome and located downstream of exon 3b (77), was predicted to increase PE inclusion (SpliceAI: 0.23) and was validated with a +0.13 PSI increase in a bulk-edited population with an average of 41% editing efficiency (Fig. 5B). We also validated two variants predicted to indirectly increase PE inclusion by disrupting canonical exon recognition, thereby favouring PE-containing isoforms. The *SLC12A6* c.2633-1G>A variant, located upstream of exon 23 had a SpliceAI score of 0.99 for loss of canonical splicing and produced a modest but reproducible increase in PE inclusion (+0.04 PSI, 21% editing efficiency; Fig. 5C). Similarly, *TSC2* c.4850-1G>A (78), was predicted to disrupt exon 33 splicing (SpliceAI: 0.99) and resulted in a +0.06 PSI change in PE inclusion (10% editing efficiency; Fig. 5D). Finally, we also investigated one variant that results in decreased PE inclusion: *COQ9* c.521+1del in (79), disrupted the canonical donor site of exon 4. SpliceAI predicted a reduction in PE inclusion (score: 0.67), which was confirmed by a –0.28 PSI change in a bulk-edited population with an average of 29% editing efficiency (Fig. 5E). Together, these results highlight the diverse ways in which pathogenic variants can impact PE splicing, either by enhancing or suppressing PE inclusion, or by disrupting canonical exon recognition.

**Figure 5.**
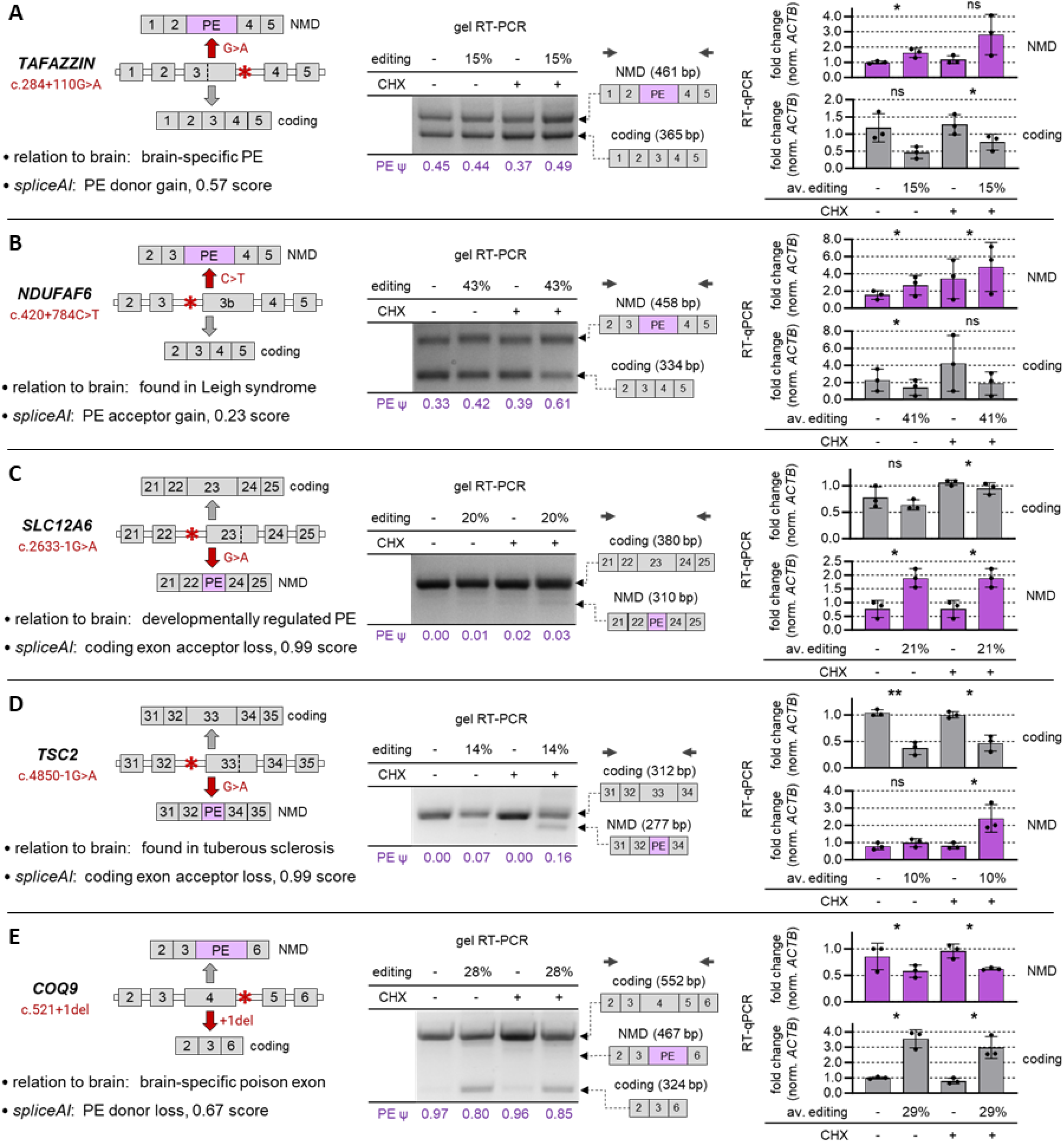
Validation of the effect of pathogenic variants on PE splicing. (**A**) c.284+110G>A, affecting E3 of *TAFFAZIN*; (**B**) c.420+784C>T, affecting E3b of *NDUFAF6*; (**C**) c.2633-1G>A, affecting E23 of *SCL12A6*; (**D**) c.4850-1G>A, affecting E33 of *TSC2*; (**E**) c.521+1del, affecting E4 of *COQ9*. The following information is reported for each variant: genomic position, disease association, spliceAI prediction, and expected RNA isoforms. Central panels show RT-PCR analysis of splicing in HEK293T cells carrying the respective variant introduced via prime editing. Cells were treated with CHX or DMSO (vehicle) 16 h prior to collection. The percentage of edited cells, measured by amplicon-NGS, is reported for each experiment. RNA isoforms with numbered exons corresponding to each PCR band are reported on the right, along with the PCR primers (black arrows). Band identity was confirmed by Sanger sequencing. PSI of the PE-containing isoform (calculated by densitometry analysis) is reported for each lane. Right panels indicate the quantitative analysis of splicing alternation by RT-qPCR. The average percentage of edited cells for each experiment. Primers specific for either the NMD isoform (purple bars) or the coding isoform (grey bars) were used. All values are internally normalized on the expression of *ACTB* gene, while transcript levels are represented as fold change compared to the DMSO condition with no editing. Statistical significance by Student’s t-test is shown for each comparison. * p<0.05, ** p<0.01, *** p<0.001, **** p<0.0001, ns: not significant. Data are shown as average ± standard deviation. Each data point represents one of three replicates.

We found of particular interest the intronic c.420+784C>T variant in the *NDUFAF6* gene, classified as “Pathogenic/Likely pathogenic” in ClinVar and associated with Leigh syndrome, a neurodevelopmental disease associated with mitochondrial complex I deficiency (77). This deep intronic mutation causes retention of 124 intronic nucleotides and creates a premature stop codon in exon 4 (77) (Fig. 5B). *NDUFAF6* has recently been shown to play a role in mitochondrial complex I assembly in HEK293T cells (80). To investigate the phenotypic consequences of *NDUFAF6* PE mis-splicing in neurons, we generated clonal i3N cell lines with different genotypes for c.420+784C>T (C/C, C/T and T/T; Fig. 6A) using the CRISPR prime editing workflow optimized in HEK293T cells. We assessed *NDUFAF6* PE splicing in undifferentiated iPSCs, immature induced neurons, and i3N neurons at 1 and 21 days of maturation. In wild-type neurons the PE is weakly included with an average PSI of 0.13 and no significant changes across differentiation stages. Heterozygous (C/T) and homozygous (T/T) mutants showed increased PE inclusion, with average PSI of 0.37 and 0.94, respectively, which remained relatively stable throughout differentiation (Fig. 6B and 6C). These results confirmed the original prediction of “acceptor gain’ for the PE (spliceAI score: 0.23). We treated the lines with CHX to assess the responsiveness of the NMD isoform (Fig. 6C). RT-qPCR analysis showed a clear increase in the PE-containing transcript upon NMD inhibition, consistent with its degradation under normal conditions. The NMD isoform was also detectable in untreated cells, in line with the well-established incomplete efficiency of NMD (67–69). To evaluate the impact of PE inclusion on NDUFAF6 protein amount, we performed Western blot analysis (Fig. 6D), which showed a significant reduction in protein levels in the homozygous mutant clone compared to wild-type cells, but not in the heterozygous. We then assessed the impact on neuronal differentiation. Using RT-qPCR, we measured the expression of pluripotency markers *NANOG*, *SOX2* and *OCT4* (81), as well as neuronal differentiation markers *VGLUT1, MAP2*, *RBFOX2* and *TUBB3* (70) (Supplementary Fig. S16A). Interestingly, homozygous (T/T) mutant cells, but not heterozygous, showed significantly higher level of *MAP2* and *RBFOX2* on day 21 of maturation (Supplementary Fig. S16A), possibly indicating an alteration in the differentiation status. Qualitative analysis by immunocytochemistry of differentiation markers MAP2, NeuN, TUJ1 and SMI-312 revealed no significant differences (Fig. 6E and Supplementary Fig. S16B). However, we observed differences in cellular organization, with both heterozygous and homozygous mutant cells forming smaller clusters. Quantitative analysis of neuronal interconnectivity at early (day 1) and late (day 21) stages of maturation confirmed this observation, showing smaller and more fragmented cell-body clusters in homozygous mutant cells. Neurite formation and branch points appeared unaffected by PE mis-splicing (Supplementary Fig. S16C and S16D). Finally, we evaluated the functional consequences of NDUFAF6 protein reduction by measuring mitochondrial membrane potential (MMP) using tetramethyl rhodamine methyl ester (TMRM) staining (82–85), which demonstrated impaired mitochondrial function in both mutant line compared to the wild type, with a more pronounced effect in homozygous mutant cells (Fig. 6F). Together, these results provide functional evidence that the c.420+784C>T variant impairs mitochondrial function, consistent with a mechanism in which poison exon inclusion reduces protein levels and leads to a disease-relevant cellular phenotype (86–89).

**Figure 6.**
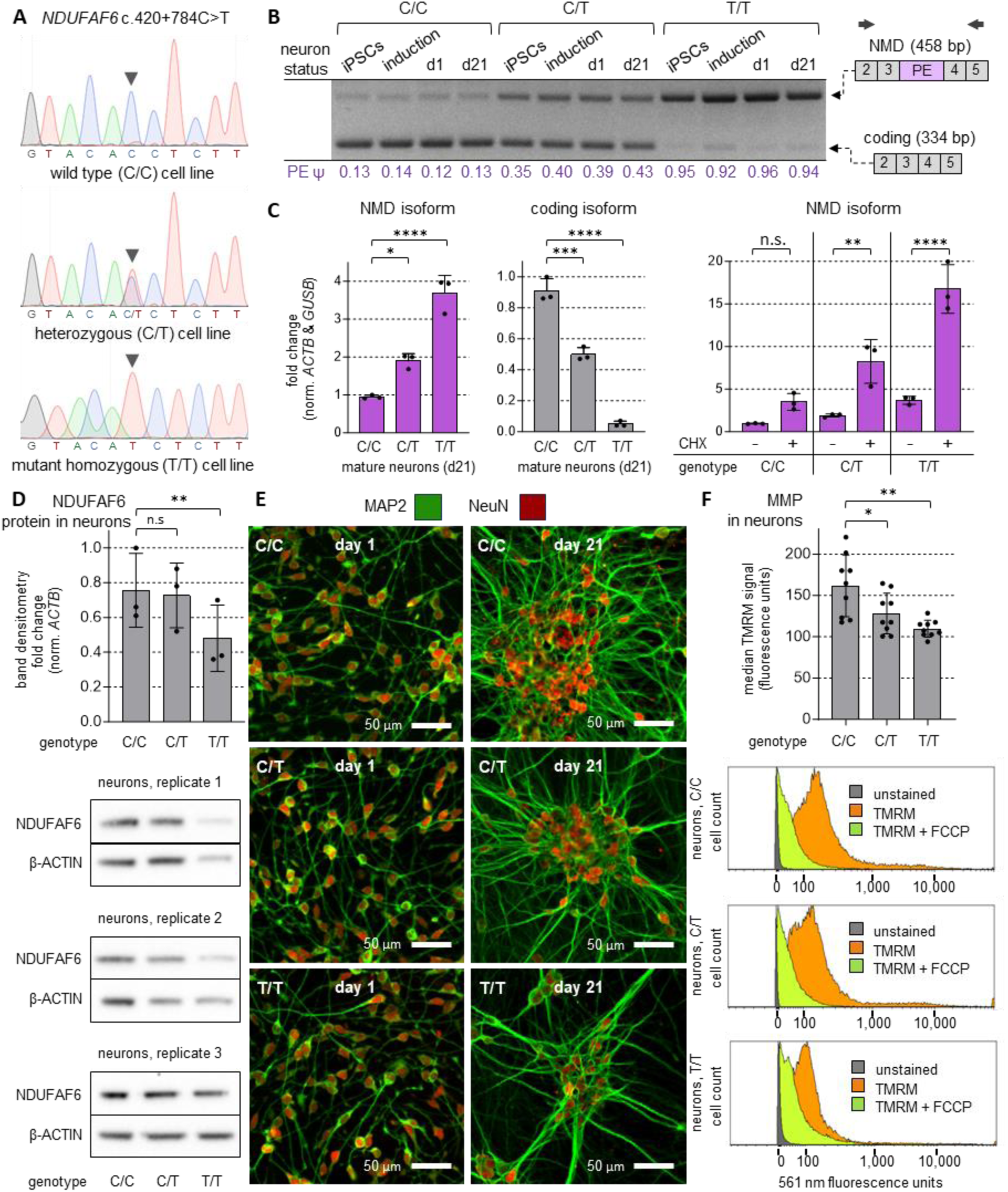
Functional characterization of *NDUFAF6* PE mis-splicing in neurons. (**A**) Genotyping by Sanger sequencing of clonal cell lines that have been prime edited to carry the mutation c.420+784C>T. Arrowheads indicate the edited base. Sequencing was performed on PCR amplicons obtained from genomic DNA. (**B**) Splicing analysis by RT-PCR of clonal cell lines with different genotypes for c.420+784C>T and at different stages of differentiation: undifferentiated iPSCs (“UN”), induced neurons following 3 days of NGN2 overexpression (“IN”), and neurons at 1 or 21 days of maturation. RNA isoforms with numbered exons corresponding to each PCR band are reported on the right, along with the PCR primers (black arrows). Band identity was confirmed by Sanger sequencing. PSI of the PE-containing isoform (calculated by band densitometry analysis) is reported for each lane. (**C**) Quantitative analysis of splicing alternation by RT-qPCR. Primers specific for either the NMD isoform (purple bars) or the coding isoform (grey bars) were used. All values are internally normalized on the expression of *ACTB* and *GUSB*, while transcript levels are represented as fold change compared to the DMSO condition with no editing. (**D**) Western blot analysis of NDUFAF6 protein expression in neurons at day 21 of differentiation and with different genotypes. Data points in the bar plot represent densitometry values from different replicates, normalized to actin and relative to the wild type (G/G). Western blots from each replicate are shown beneath the plot. (**E**) Immunocytochemistry analysis of c.420+784C>T clonal cell lines, at 1 or 21 days of maturation, using antibodies for neuronal markers MAP2 and NeuN. Images are representative of multiple fields and replicates. (**F**) MMP is measured in neurons at day 21 of differentiation and with different genotypes by TMRM staining. Fluorescence was detected using a flow cytometer with 561 nm laser and a 548/574 nm emission filter. Data points in the bar plot represent median fluorescence of the cell population from different replicates. The fluorescence profile of the cell population is show beneath the plot for selected replicates of each genotype, either unstained, stained with TMRM, or stained with TMRM and treated with 10 µM FCCP. In (C, D, F), statistical significance by one-way ANOVA with Geisser-Greenhouse correction is shown for each comparison. * p<0.05, ** p<0.01, *** p<0.001, **** p<0.0001, ns: not significant. Data are shown as average ± standard deviation. Each data point represents one of multiple replicates.

Overall, these findings establish PE dysregulation as a widespread, clinically relevant mechanism in the nervous system and highlight its potential as a therapeutic target.

## Discussion

Alternative splicing coupled with NMD is a central mechanism for regulating transcript stability and fine-tuning gene expression in eukaryotes. PE inclusion represents a specialized, highly conserved splicing event that triggers NMD and reduces protein expression. Although isolated examples of PEs have been described, their broader role in human brain development and disease remains poorly understood. Here, we provide a comprehensive characterization of the role of PE in human gene expression regulation by integrating evolutionary, transcriptomic, and disease-linked datasets with functional validations. Our study identified thousands of human PEs with tissue- and developmentally regulated splicing patterns in the brain, as well as hundreds of pathogenic variants affecting their splicing.

The comparative analysis of 957 eukaryotic transcriptomes revealed that NMD isoforms are enriched in vertebrates, especially in *Homo sapiens*. We estimate that 39.6% of human coding genes give rise to at least one NMD transcript (30.3% if considering only transcripts with TSL 1-3), and 20.1% of all protein-coding transcripts are predicted to undergo NMD (14.3% if considering only transcripts with TSL 1-3). Together, these results suggest that NMD plays a prominent role in shaping transcriptome composition and regulating gene expression in *Homo sapiens*. Species with more complex splicing regulation have evolved to utilize NMD for fine-tuning gene expression at the post-transcriptional level. Within this context, PEs likely represent an evolutionarily refined strategy to control gene dosage in a spatial-temporal manner, especially in complex organs such as the brain. These analyses are based on annotation-driven predictions and should be interpreted carefully, as NMD rules may differ across species (90, 91) and annotation depth varies, with more comprehensively annotated genomes (such as *Homo sapiens*) potentially overrepresenting low-abundance alternative isoforms and thus influencing the predicted frequency of PEs.

We mapped the PE landscape, and we identified over 9,000 putative PEs across the human transcriptome, accounting for nearly 18% of all annotated alternative splicing events. These PEs were enriched in genes involved in RNA metabolism and in splicing factors, some of which are known to autoregulate their own expression through negative feedback loops mediated by PE inclusion (19, 27, 59). While longer genes were more likely to contain PEs, those with brain-specific expression showed a significant depletion of PEs, suggesting selective pressure to preserve expression of critical neural genes. Experimental validation confirmed that PE inclusion typically leads to NMD, validating our predictive framework. The overlap of PE-containing genes with neurological disorder–associated pathways underscores the clinical significance of PE splicing as a key regulatory mechanism in the human brain.

We next investigated PE regulation in the context of the human brain. Using RNA-seq data from GTEx, we identified PEs that are differentially spliced in the brain compared to other human tissues. Analysis of BrainSpan RNA-seq data revealed dynamic changes in PE splicing across brain development, with distinct shifts occurring before and after birth. We observed significant changes in PE inclusion during neuronal differentiation, as for example in *NOVA1*, suggesting tight regulation of PE inclusion during neurogenesis. Together, our findings indicate that PEs act as temporally and spatially regulated molecular switches that fine-tune gene expression in both the developing and adult brain.

To assess the clinical impact of PE mis-splicing, we integrated ClinVar data with SpliceAI predictions and identified pathogenic variants predicted to affect PE splicing in neurological disorders. Importantly, ClinVar classification of variants as “pathogenic” or “likely pathogenic” does not necessarily imply that the underlying molecular mechanism has been fully characterized. In particular, many of these variants are located in intronic regions, where pathogenicity is often supported by genetic or clinical evidence, while the precise splicing defect remains unknown. In this context, our approach provides a mechanistic layer of interpretation by identifying variants whose predicted effects are consistent with PE mis-splicing, an underexplored regulatory mechanism. This study therefore enables a more refined functional annotation of disease-associated variants and helps uncover previously unrecognized splicing mechanisms that may contribute to human disease. Using CRISPR prime editing (49), we functionally validated representative variants selected based on their involvement in relevant neurological disorders and their effect on splicing, and confirmed that they induced alterations in endogenous mRNA splicing in human cells validating the accuracy of our predictions. Since SpliceAI has known limitations in predicting aberrant PE inclusion, particularly for weakly spliced or non-annotated exons (44), our pipeline focused on annotated exons which generally exhibit stronger splice site signals and therefore support more reliable predictions. Although previously recommended thresholds include Δ score >0.1 for splice loss and >0.5 for splice gain (2), we adopted a uniform cutoff of 0.2 while retaining the full SpliceAI scores to enable flexible downstream filtering depending on the desired stringency. We identified variants associated with a broad spectrum of neurological conditions, including epilepsy, intellectual disability, and neurodegeneration, highlighting the role of PE mis-splicing in brain disorders. We further examined the consequences of the *NDUFAF6* c.420+784C>T variant, associated with Leigh syndrome, in the context of neuronal cells. After generating heterozygous and homozygous mutant i3N neurons, we observed an allele-dependent increase of the inclusion of a PE, confirming our original computational prediction. We then functionally characterized the consequences of PE mis-splicing on NDUFAF6 protein levels and mitochondrial function, establishing a direct link between this splicing defect and the observed cellular phenotype. Homozygous mutant neurons displayed near-complete PE inclusion, a marked reduction in NDUFAF6 protein expression, and impaired mitochondrial membrane potential, while heterozygous cells showed intermediate phenotypes. The observed disruption in mitochondrial membrane potential is consistent with the established role of NDUFAF6 in mitochondrial complex I assembly (77, 92), further supporting the pathogenic relevance of PE mis-splicing in this context. To our knowledge, this represents the first functional characterization of NDUFAF6 deficiency resulting from PE mis-splicing in a human neuronal model. The phenotypic severity was substantially more pronounced in homozygous mutant cells, consistent with clinical observations that most patients carrying pathogenic *NDUFAF6* variants are either homozygous or compound heterozygous for deleterious alleles(77, 86–89, 93, 94). We additionally observed altered expression of the neuronal differentiation markers *MAP2* and *RBFOX2* in the homozygous mutant cells. Of note, the identification and characterization of this PE is particularly relevant from a therapeutic perspective, as NMD-sensitive PE events represent potentially targetable splicing defects that could be corrected using emerging RNA-based therapeutic approaches (95). Overall, our experimental pipeline provides a scalable and adaptable framework for interpreting and validating splicing variants affecting PE splicing within their native transcriptomic context.

## Conclusions

This study establishes PEs as a widespread, evolutionarily conserved, and dynamically regulated mechanism of gene expression regulation in the human brain. By integrating computational prediction with functional validation, we show that PEs are critical regulators of gene expression during brain development and that their mis-splicing constitutes a recurrent pathogenetic mechanism in neurological disease. These findings highlight the therapeutic potential of targeting PE mis-splicing and provide a foundational resource for advancing the study of splicing regulation in the human brain and its contribution to disease.

## Methods

### Evolutionary analysis

Genome annotations of Eukaryotic species were downloaded in GTF format from Ensembl Genomes Release 60 (https://ftp.ebi.ac.uk/ensemblgenomes/pub/release-60/) and Ensemble Release 113 (for vertebrates, https://ftp.ensembl.org/pub/release-113/) using a Python-based script. Species are classified in this study based on their original group classification in the repository, which includes “vertebrates”, “metazoan” (here indicated as “invertebrates”), “protists”, “fungi” and “plants”.

A Python-based script was developed to systematically analyse each genome annotation for predicting NMD isoforms and collecting genomic information as follows: 1) unique isoforms are identified by their “transcript_id”, while coordinates of “exon” and “CDS” features are collected and annotated for each isoform; 2) the relative position of the last CDS coordinate (indicating the annotated stop codon) is analysed in relation to the position of the last exon-exon junction; 3) isoforms for which the stop codon is located more than 50 nt upstream of the last exon-exon junction are annotated as “NMD”; 4) additional information is collected from each genome annotation, including i) the total number of coding genes (uniquely identified by their “gene_id”, and for which at least one isoform with a “CDS” feature has been annotated), ii) the number of unique isoforms per coding gene or in total, iii) the number of unique “exon” feature per coding gene, iiii) the length of each coding isoform.

For phylogenetic tree reconstruction, taxonomic IDs of each species and lineage information were both retrieved from NCBI public repositories (respectively https://ftp.ncbi.nih.gov/genomes/ASSEMBLY_REPORTS/assembly_summary_refseq.txt and https://ftp.ncbi.nih.gov/pub/taxonomy/taxdump_archive/new_taxdump_2023-12-01.zip). The phylogenetic tree was generated with the ETE package (96), using the function ncbi.get_topology(taxids, intermediate_nodes=True) and lineage information from the file rankedlineage.dmp (retrieved in new_taxdump_2023-12-01.zip). The tree illustration was generated using iTOL online tool (97).

To assess annotation quality across species, transcriptome completeness was evaluated using BUSCO v5 (50) in transcriptome mode with the universal eukaryotic lineage dataset “eukaryota_odb12”. Highly complete annotations were defined as those with BUSCO completeness scores ≥90%. Additional analyses for *Homo sapiens* and *Mus musculus* were performed using only transcripts annotated with Ensembl transcript support levels (TSL) 1–3, corresponding to isoforms supported by stronger experimental evidence.

The original annotations of *Drosophila melanogaster* and *Arabidopsis thaliana*, which did not contain transcripts explicitly annotated with the “nonsense_mediated_decay” biotype, were implemented using a custom Python-based procedure to identify and annotate putative NMD-sensitive isoforms. The script parses transcript annotations from the original GTF file, and transcripts predicted to undergo NMD according to the pipeline described above are reannotated by assigning the “nonsense_mediated_decay” transcript biotype.

All codes and procedures are described in the Supplementary Methods and are available at the following GitHub repository: https://github.com/ppigini/EvoNMD.

### Identification of putative PEs from genomic and transcriptomic reference

This study focuses on defining the PE landscape by leveraging the annotated genome (GRCh38) and transcriptome (Gencode v26), based on the assumption that an annotated exon can be alternatively included as a PE in an otherwise protein-coding transcript and result in a downstream PTC. The derived PTC is not necessarily within the PE itself but has to be more than 150 bp from the transcription start site and not in the final two exons of the host gene, ensuring it doesn’t escape NMD (51). To search for PE from the transcriptome reference, non-redundant exons whose length ≤ 300 bp of a gene were first extracted from the GTF file. Each extracted exon was then computationally inserted into each transcript of the gene whose “transcript_biotype” is “protein-coding”. If the insertion generated a PTC meeting the criteria above, the inserted exon would be marked as a putative PE. Note, this also means that whether an exon is “poisoning” or not depends on its transcript context. Depending on the relative position of the putative PE and the protein-coding transcript, the PE can be classified as one of “inclusion”, “alternative 5’-ss” and “alternative 3’-ss” (Supplementary Fig. S3A).

### Brain-expressed genes and brain-unique splice junction

Brain-expressed genes were defined using median transcripts-per-million (TPM) values from the GTEx v8 dataset. For each gene, the median TPM was calculated across all brain regions and across all non-brain tissues separately, as well as the maximum TPM observed in any single non-brain tissue. A gene was classified as brain-expressed if it met either of two criteria: 1) it had zero or negligible expression in non-brain tissues (maximum non-brain TPM ≤ 1), or 2) its median brain TPM was more than 50-fold greater than its median non-brain TPM. Using these criteria, 1,745 brain-expressed genes were identified. Brain-unique splice junctions were identified from GTEx junction-level read counts. For each junction, read counts were summed across all samples within each tissue, and the median read count was calculated across all brain tissues and all non-brain tissues separately. A junction was classified as brain-unique if it was detectably expressed in brain tissues (median brain read count > 2) and showed strong brain enrichment (ratio of median brain to median non-brain read count > 50). Using these criteria, 1,092 brain-expressed junctions were identified.

To assess whether brain-expressed genes were enriched among genes containing increasing numbers of PEs, we performed one-tailed hypergeometric tests for each PE-count bin. The full set of protein-coding genes was used as the background population, and p-values were corrected for multiple testing using the Benjamini–Hochberg FDR method (FDR < 0.05).

### PSI value calculation for PEs

To quantify the inclusion level of each PE across tissues and developmental stages, we computed the PSI value based on aligned read coverage from short-read RNA-seq data obtained from GTEx and BrainSpan. For each PE event, the read counts mapped to the alternative exon region were extracted and denoted as *R₁*, *R₂*, and *R₃*, respectively. The PSI value was then calculated based on different PE modes. For PE inclusion, the PSI value was calculated as:

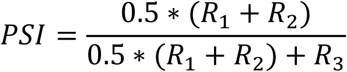

For alternative 5’/3’-ss mode, PSI was calculated as:

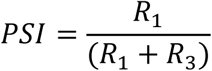

Note, this should not be confused with the inclusion of an exon being explicitly used as a PE. From the short-read data used in this study, only the overall inclusion of an exon is measurable, regardless of whether it is included in a transcript as a poison or non-PE.

### Identification of tissue-specific PEs from GTEx

The sample-wise PSI profiles were first built for all GTEx samples and then the PSI values were logit-transformed with the exactly 0 replaced by 0.001 and values of exactly 1 replaced by 0.999 prior to applying the logit transformation. For every pair of tissues, a generalized linear model (GLM) was fitted to examine the relationship between PSI and tissue type for every splicing event involving the exon with PE potential, the model was:

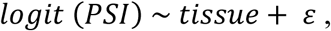

With tissue as a two-level factor. Sex, age group and ancestry were not included as confounding variables as they showed no systematic separation along the leading principal components (Supplementary Fig. S4A-C). The direction and significance of the coefficient for tissue type were used to judge whether the exon is differentially used between brain tissue and other tissues.

### Identification of developmental-time-specific PEs from BrainSpan

To identify PEs whose inclusion changes significantly across human brain development, we modelled PSI values as a function of developmental stage using Generalized Additive Models (GAMs) for each brain region respectively. PSI values at the boundaries were capped by replacing values of exactly 0 with 0.001 and values of exactly 1 with 0.999 prior to applying the logit transformation. PSI values strictly between 0 and 1 were not modified. The capped PSI values were then logit-transformed and used as the response variable in all downstream models. For each PE in each brain region, the logit-transformed PSI was modelled as:

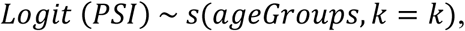

with developmental age group modelled as a smooth term with default parameters, and a Gaussian family with identity link. PCA of PSI matrices across brain regions showed no systematic separation by sex (Supplementary Fig. S6) and ancestry (Supplementary Fig. S7), so they were not included in the models. PEs with significant developmental dynamics were defined as those with smooth-term p < 0.05 after FDR correction.

To group PEs with similar developmental trajectories, unsupervised hierarchical clustering was performed on the GAM-fitted logit(PSI) values of all significant PEs using Pearson correlation distance. The resulting dendrogram was cut at a fixed height of 0.2, and the clusters containing the highest number of PEs were retained to focus on dominant trajectory patterns. For each retained cluster, PCA was performed on the matrix of fitted trajectories and extracted the first principal component (PC1) score across samples. This yields a sample-length vector that summarizes the dominant pattern of variation in logit(PSI) across development among PEs in the cluster. To ensure that the direction of the PC1 trajectory matched the dominant biological trend rather than its arbitrary numerical inverse, the sign of PC1 was flipped when its correlation with the cluster’s mean trajectory was negative. The resulting PC1 trajectories were used for downstream visualization and cross-region comparisons.

### Analysis of pathogenic variants

Pathogenic and likely-pathogenic mutations were extracted from ClinVar VCF file (version 20231112). SpliceAI (2) or Pangolin (45) were applied to predict the effect on splicing for each mutation. A Python-based workflow (https://github.com/ppigini/SpliceDirect) was developed to filter the variants analysed with SpliceAI or Pangolin, match them with PE inclusion data, and identify those affecting PE splicing. Only the most impacted putative splice site (ss) with an absolute SpliceAI score change > 0.2 were retained. The genomic coordinates of these impacted ss were then matched with the coordinates of the PE’s ss. When an exact match was found, the mutation would be considered to affect the splicing of the PE. Then, for each variant, we additionally assigned a positive or negative value to indicate directionality, where positive scores indicate increased PE inclusion, and negative scores indicate decreased inclusion (significant absolute value > 0.2). We paired these splicing-change predictions with the wild-type PSI values from GTEx brain tissues to prioritize variants that significantly affect PE splicing in the brain. The following two scenarios were of interest: 1) a PE’s PSI in wild-type is < 0.3 and the mutation is predicted to increase its inclusion; and 2) a PE’s PSI in wild-type is > 0.7 and the mutation is predicted to decrease its inclusion. Variants and genes associated with brain disorders were identified by looking for specific disease terminology in the ClinVar report (if available) or in the GO respectively. Gene ontologies were obtained using the Metascape online tool (98). Specific terminology for brain disorders was retrieved from the ODiseA database (99). All codes and procedures are described in the Supplementary Methods and are available at the following GitHub repository: https://github.com/ppigini/SpliceDirect.

### Cell culturing

HEK293T cells (ATCC CRL-3216) were maintained in Dulbecco’s Modified Eagle Medium (DMEM, Gibco™ 11965118) supplemented with 10% fetal bovine serum (FBS, HyClone™ SH3008003) and 1% penicillin-streptomycin (Gibco™ 15140122) at 37°C in a humidified incubator with 5% CO₂.

Human iPSCs carrying inducible NGN2 (Coriell Institute GM29370) were differentiated into i3N neurons following a multi-step protocol adapted from Wang et al. (2018) (70). Plates were coated with Matrigel (Corning™ 354277) diluted in cold DMEM/F12 (Gibco™ 11320082) as per manufacturer’s instructions, and incubated at room temperature for at least 3 hours. On day -3, iPSCs at ∼80% confluency and cultured in mTeSR Plus™ medium (StemCell™ 100-0276) with 1% penicillin-streptomycin (Gibco™ 15140122) were dissociated with Accutase™ (StemCell™ 7920) and plated on Matrigel-coated dishes in induction medium, which was composed of KO-DMEM/F12 (Gibco™ 12660012) supplemented with N2 (Gibco™ 17502048), NEAA (Gibco™ 11140050), 10 ng/mL BDNF (PeproTech™ 450-02), 10 ng/mL NT3 (PeproTech™ 450-02), 0.2 µg/mL laminin (Corning™ 354239), 2 µg/mL doxycycline (Sigma-Aldrich™ D9891-1G), and 10 µM ROCK inhibitor (Sigma-Aldrich™ 688001). Medium was changed with complete induction medium on day - 2, and with induction medium lacking ROCK inhibitor on day -1. On day 0, cells were dissociated, replated onto dishes coated with 50 µg/mL poly-D-lysine (Gibco™ A3890401) and 3 µg/mL laminin (Gibco™ 23017015), and cultured in maturation medium, which was composed of 50% DMEM/F12 (Gibco™ 11320082) and 50% Neurobasal-A™ (ThermoFisher™ 10888022), supplemented with 0.5× B27 (Gibco™ 17504044), N2 (Gibco™ 17502048), NEAA (Gibco™ 11140050), GlutaMax (Gibco™ 35050061), 1 µg/mL laminin (Corning™ 354239), 10 ng/mL BDNF (PeproTech™ 450-02), and 10 ng/mL NT3 (PeproTech™ 450-02). Half-medium changes were performed weekly starting from day 7. Neurons were cultured for up to 4 weeks before downstream assays. Cells were cultured at 37°C in a humidified incubator with 5% CO₂ at all stages. Live-cell imaging was performed using a Sartorius Incucyte SX5 system (Sartorius™) equipped with a 10× objective, and images were analysed using the integrated NeuroTrack software.

CHX treatment was performed both on HEK293T and i3N neurons by treating them for 16h with 50 μg/ml CHX (Thermo Scientific™ J66901.03) or an equivalent volume of DMSO (Sigma-Aldrich™ D8418) as control (0.5% of the culture medium volume). For HEK293T cells, medium was replaced with CHX-containing medium. For i3N neurons, culture medium was quickly removed from the cells, implemented with CHX, and placed back in culture.

### CRISPR Prime Editing

Plasmids expressing the CRISPR prime editor (Addgene 174820), the pegRNA (Addgene 223137), or the nicking gRNA (Addgene 65777) were prepared using competent DH5α E. coli (NEB™ C2987I) and QIAwave Plasmid Miniprep Kit (Qiagen™ 27204). pegRNAs and nicking gRNAs (Supplementary Table S12) were designed using the PRIDICT tool (75) and selecting the pegRNA and gRNA with top scores, and they were cloned using standard Type IIS restriction enzymes (NEB™). Sequences were confirmed by Sanger sequencing.

For CRISPR prime editing of HEK293T, cells were seeded at 100,000 per well in 24-well plates one day prior. Transfections were performed using Lipofectamine 3000™ (Invitrogen™ L3000008) according to the manufacturer’s protocol. Briefly, DNA plasmids and P3000 reagent™ were diluted in Opti-MEM™ (Gibco™ 31985070), combined with Lipofectamine 3000™ diluted in Opti-MEM™, incubated for 15 minutes at room temperature, and added dropwise to the cells in DMEM with FBS but no antibiotics. Medium was replaced after 8 hours and doubled after 24 hours. Cells were collected 72 hours post-transfection for downstream analyses.

For CRISPR prime editing of iPSCs, cells were transfected with plasmids expressing the editing components using Lipofectamine™ Stem Transfection Reagent (Invitrogen™, STEM00001). GFP-positive cells were FACS sorted as single cells in 96-well plates using BigFoot™ Cell Sorter (Thermo Fisher™) to generate isogenic lines. Clonal lines were cultured for about 2 weeks before splitting them for genotyping by Sanger sequencing and downstream freezing. Cells were frozen using a solution of 50% DMEM/F12 (Gibco™ 11320082) and 50% heat-inactivated FBS (CellPro™ FB73), supplemented with 10% DMSO (Sigma-Aldrich™ D8418).

Genome editing efficiency was assessed by amplicon-NGS. Genomic DNA was extracted from cultured cells using magnetic purification beads (Illumina™ 20060057) and following the manufacturer’s specifications. Briefly, cells were lysed using the provided solution, and the lysate was incubated with purification beads to bind DNA. Magnetic separation was used to wash away impurities, and DNA was eluted in nuclease-free water. To quantify editing outcomes, targeted regions were amplified from genomic DNA using primers with overhangs compatible with Illumina adapters (Supplementary Table S12). Amplicons were cleaned with magnetic beads (Illumina™ 20060057) and pooled for multiplexed sequencing. Libraries were sequenced with MiSeq Reagent Micro Kit v2™ (Illumina™ MS-103-1002) on a MiSeq Syste™ (Illumina™) using 2×150 bp paired-end reads. Data were demultiplexed and analyzed using the CRISPResso2 tool (100).

### *In vitro* splicing analysis

Total RNA was extracted from cultured cells using QIAzol (Qiagen™ 79306) reagent according to the manufacturer’s instructions. After chloroform separation (Sigma-Aldrich™ C2432) and isopropanol precipitation (Sigma-Aldrich™ 109634), RNA pellets were washed in 75% ethanol (Sigma-Aldrich™ E7023), air-dried, and resuspended in nuclease-free water. RNA quantity and purity were assessed using a NanoDrop spectrophotometer (Thermo Scientific™ 2353-30-0010). For reverse transcription, 500 ng to 1 µg of total RNA was used with the SuperScript III Kit (Fisher Scientific™ 18080044), using random hexamer primers (Promega™ C1181) and following the recommended protocol.

PCR was performed using exon-specific primers (Supplementary Table S12) and GoTaq master mix (Promega™ M7123-Green) to amplify regions of interest from cDNA (25 µg per reaction). PCR products were resolved on 2% agarose gels (Fisher Scientific™ 16-550-100) stained with ethidium bromide (Sigma-Aldrich™ E1510) and visualized using a UV gel imaging system. Bands of expected size were excised and purified using the QIAquick Gel Extraction Kit (Qiagen™ 28706), followed by Sanger sequencing to confirm splicing isoforms and identify sequence variants. Images of gel bands were acquired using AlphaImager™ 2200, and densitometry analysis was performed in ImageJ (101).

qPCR was carried out using SYBR Green PCR Master Mix™ (Fisher Scientific™ A25742) in a LightCycler 480 II system™ (Roche™). Reactions were performed in technical duplicates using 10 µg cDNA per reaction and exon-specific or exon junction-specific primers (Supplementary Table S12). Relative gene expression was calculated using the ΔΔCt method (102), normalized to housekeeping genes. Melting curve analysis was used to ensure specificity and consistency across runs.

### Sashimi plot visualization

To provide orthogonal evidence to qPCR measurements of splice junction for the PE inclusion events, RNA-seq from public available HEK293T (GSE239877) and home-made i3N were used to generate Sashimi plots using the Gviz package for the key PE events shown in Fig. 5 and Fig. 6.

### Western blot analysis

Protein extracts were prepared from iPSCs and day 21 neurons using RIPA lysis buffer supplemented with DTT, SDS, protease/phosphatase inhibitors, and Benzonase. Cell pellets were resuspended in lysis buffer, subjected to three freeze–thaw cycles, and centrifuged at 13,000 × g for 10 min at 4°C to collect the soluble protein fraction. Protein concentration was determined using the Pierce BCA Protein Assay Kit (Thermo Fisher Scientific). Equal amounts of protein (30 μg) were mixed with Laemmli sample buffer containing β-mercaptoethanol, denatured at 95°C for 5 min, and separated on 4–12% Bis-Tris polyacrylamide gels (NuPAGE, Thermo Fisher Scientific). Proteins were transferred onto PVDF membranes and blocked in 5% milk in TBST for 1 h at room temperature. Membranes were incubated overnight at 4°C with primary antibodies against NDUFAF6 (Biorbyt ORB3068405, 1:500) and β-Actin (Cell Signaling Technology 4970T, 1:10,000), followed by HRP-conjugated anti-rabbit secondary antibody (Cell Signaling Technology 7074P2, 1:5,000). Chemiluminescent detection was performed using SuperSignal West Pico PLUS or SuperSignal West Femto substrates (Thermo Fisher Scientific), and signal acquisition was performed using the iBright imaging system (Thermo Fisher Scientific).

### Immunocytochemistry

Cells cultured in 96-well plates (Corning™ 3603) were fixed by adding an equal volume of 8% paraformaldehyde (Thermo Fisher™ 047347.9M) in PBS (Gibco™ 10010023) directly to the culture medium (final concentration 4% paraformaldehyde) and incubating for 30 min at room temperature. After fixation, cells were washed three times with DPBS without calcium and magnesium (Gibco™, 14190114), taking care to avoid drying, and permeabilized with 0.1% Triton X-100 (Thermo Scientific™ PI85111) in DPBS for 10 min at room temperature. Following three additional DPBS washes, cells were blocked in DPBS containing 5% normal goat serum (Sigma-Aldrich™ S26-100ML) and 0.01% Triton X-100 for 1 h at room temperature on a shaker. Primary antibodies diluted in blocking solution were applied overnight at 4 °C (see antibody details below). The following day, cells were washed three times with DPBS and incubated for 1 h at room temperature with the corresponding secondary antibodies diluted in blocking solution, protected from light. After washing, nuclei were counterstained with 2 µg/mL Hoechst-33342 (BioRad™ 1351304EDU) for 10 min, followed by three DPBS washes. Wells were finally filled with DPBS for imaging. The following primary antibodies were used: NeuN (Millipore™ MAB377, clone 60), MAP2 (Millipore™ AB5543), βIII-tubulin (Synaptic Systems™ 302304), PSD95 (NeuroMab™ 75-028), and SMI-312 (Abcam™ ab254348). Secondary antibodies were: goat anti-mouse Alexa Fluor™ 594 (Thermo Fisher™ A11032), goat anti-chicken Alexa Fluor™ 488 (Thermo Fisher, A11039), goat anti-guinea pig Alexa Fluor™ 488 (Thermo Fisher™ A11073), and goat anti-rabbit Alexa Fluor™ 594 (Thermo Fisher™ A11012). Images were acquired using an IN Cell Analyzer 6000 (Cytiva™) with a 20× objective, capturing 30 fields per well, and analysed in ImageJ (101) for channel assignment, brightness/contrast adjustment.

### TMRM assay

Mitochondrial membrane potential was assessed in day 21 neurons using tetramethylrhodamine methyl ester (TMRM; Invitrogen I34361). Cells were incubated for 30 min at 37°C in maturation medium lacking BDNF and NT3 and containing 25 nM TMRM. As a positive control for mitochondrial depolarization, cells were treated with 25 nM TMRM together with 10 μM FCCP (MedChemExpress HY-100410). Following incubation, cells were detached using Dispase, resuspended in FACS buffer (PBS containing 5% FBS and 1 mM EDTA), filtered through 70 μm cell strainers, and analyzed using a Bigfoot Cell Sorter (Thermo Fisher Scientific). TMRM fluorescence was detected using a 561 nm laser and a 548/574 nm emission filter. Median fluorescence intensity values were quantified from live-cell populations and analyzed using FCSalyzer software.

### Statistical analysis

The statistical analysis for tissue-wise and developmental stage PE identification were implemented using R (v 4.4.1) with GAM performed with mgcv (v 1.9.4) package and GLM performed with glm function in base R. The brain-unique PEs were defined as those with a minimal PSI > 0 in any brain tissue and a max PSI = 0 across all non-brain tissues. The brain-differential PEs were defined as those meeting all the following criteria when comparing a brain tissue to a non-brain tissue: 1) tissue coefficient ≥ 0.2 or ≤ -0.2, and 2) tissue coefficient’s false discovery rate (FDR) < 0.1 from the Wald test of GLM. The developmental-stage-specific PEs were defined as those whose smooth term’s FDR < 0.05 in the GAM. For the GO enrichment analyses, a threshold of FDR < 0.1 from Fisher’s exact test was applied to select the significant terms. The statistical analyses for the evolutionary analysis, splicing variant predictions and wet-lab work were performed using GraphPad Prism. Statistical significance by non-parametric Kruskal–Wallis test, followed by Dunn’s multiple comparisons test, was assessed for the evolutionary analysis in Supplementary Fig. S1A-C, and the SpliceAI analysis of intronic variants in Supplementary Fig. S12. Spearman correlation analysis was performed for the evolutionary analysis in Supplementary Fig. S2, the comparison between HEK293T and i3N in Supplementary Fig. S10A, and the comparison between SpliceAI and Pangolin in Supplementary Fig. S13. Statistical significance was assessed by Student’s t-test for the CHX-response analysis in Fig. 2B and Supplementary Fig. S3B and S11A-B, and for RT-qPCR results in Fig. 5 and Supplementary Fig. S14D. Statistical significance was assessed by one-way ANOVA with Geisser-Greenhouse correction for experimental characterizations in Fig. 5 and Supplementary Fig. S16. In all analyses performed with Student’s t-test or ANOVA test, statistical significance is indicated as follows: “****” indicates P < 0.0001; “***” indicates P < 0.001; “**” indicates P < 0.01; “*” indicates P < 0.05; “ns” indicates non significance.

## Supplementary information

**Additional file 1: Supplementary Figures.pdf.** Fig S1. Analysis of annotated NMD isoforms across different taxonomic groups. Fig S2. Analysis of NMD isoforms frequency across species. Fig S3. Analysis of PE splicing events predicted in the human genome. Fig S4. PCA plot of demographic covariates on PSI matrix across GTEx tissues. Fig S5. Correlation between PE inclusion and gene expression. Fig S6. PCA plot of PE inclusion across BrainSpan samples colored by Sex. Fig S7. PCA plot of PE inclusion across BrainSpan samples colored by ancestry. Fig S8. PCA plot of PE inclusion across BrainSpan samples colored by developmental age. Fig S9. PE inclusion clustered into trajectories in different brain regions. Fig S10. Analysis of i3N neurons as model for brain-specific PEs. Fig S11. Analysis of CHX-responsiveness of selected PEs in i3N neurons. Fig S12. SpliceAI analysis of 236,304 ClinVar genetic variants. Fig S13. Comparison between SpliceAI and Pangolin in predicting splicing variants. Fig S14. Validation of the effect of genetic variants on PE splicing in HEK293T cells. Fig S15. Sashimi plots for selected PEs affected by pathogenic splicing variants. Fig S16. Characterization of NDUFAF6 PE mis-splicing in i3N neurons.

**Additional file 2: Supplementary Tables.xlsx.** Table S1. Annotated NMD isoforms and genomic features for Eukaryotic species. Table S2. Alternative splicing events and putative PEs in Homo sapiens. Table S3. Alternatively spliced PEs in model organisms. Table S4. Enrichment analysis in KEGG pathways for PE-containing genes. Table S5. Enrichment analysis in UniProt KW diseases for PE-containing genes. Table S6. Brain-unique and brain-differential PEs based on GTEx expression data. Table S7. Top GO categories of genes with brain-specific PE exclusion or inclusion. Table S8. Differentially spliced PEs during brain development based on BrainSpan expression data. Table S9. GO enrichment for PE host genes from high PE inclusion rate before birth and after birth patterns. Table S10. PC1 trajectories of clustered PEs across BrainSpan developmental stages. Table S11. Pathogenic and likely-pathogenic ClinVar variants predicted to affect the splicing of brain-relevant PEs. Table S12. PCR primers and pegRNA/gRNA sequences.

**Additional file 3: Supplementary Methods.pdf.**

## Declarations

### Ethics approval and consent to participate

Not applicable.

### Consent for publication

Not applicable.

### Data and code availability

The RNA-seq for HEK293T cell line from public available GEO dataset GSE239877. The data supporting the conclusions of this article are included within the article (and its additional files), and are available in Zenodo at DOI 10.5281/zenodo.17041770. All original code has been deposited at the following GitHub repositories and is publicly available as of the date of publication:

https://github.com/ppigini/EvoNMD

https://github.com/dgaolab/Poison_Exon_Landscape

https://github.com/ppigini/SpliceDirect

### Author contributions

Paolo Pigini: Conceptualization, Data curation, Formal analysis, Investigation, Methodology, Software, Supervision, Validation, Visualization, Writing - original draft, Writing - review & editing. Huilin Xu: Conceptualization, Data curation, Formal analysis, Investigation, Methodology, Software, Supervision, Validation, Visualization, Writing - original draft, Writing - review & editing. Yan Ji: Investigation, Methodology. Hannah Lindmeier: Investigation, Methodology. Henry R. Saltzman: Investigation, Methodology. Henry Shull: Investigation, Methodology. Shuqi Yun: Investigation, Methodology. Christiano R. R. Alves: Methodology, Resources. Melissa A. Walker: Investigation, Methodology, Resources. M. Catarina Silva: Conceptualization, Funding acquisition, Methodology, Resources, Supervision. Dadi Gao: Conceptualization, Data curation, Formal analysis, Funding acquisition, Methodology, Resources, Software, Supervision, Validation, Visualization, Writing - original draft, Writing - review & editing. Elisabetta Morini: Conceptualization, Funding acquisition, Methodology, Project administration, Resources, Supervision, Validation, Writing - original draft, Writing - review & editing.

## Acknowledgements

We would like to express our sincere gratitude to Drs. Jacob Kitzman and Jennifer Yee from the University of Michigan Medical School for their invaluable initial contributions in identifying the optimal approach and models to study genetic mutations affecting RNA splicing. Their foundational work was instrumental in shaping the direction of this research. We also extend our sincere appreciation to Dr. Xander Nuttle, also from the University of Michigan Medical School, for his expert guidance in the editing of iPSCs and the subsequent selection for neuronal differentiation. His insights were crucial for the successful execution of key experimental procedures.

