## Supplemental Figures (S1-S16) for "Poison exon splicing in the human brain: a new paradigm for understanding and targeting neurological disorders"

### Additional file 1

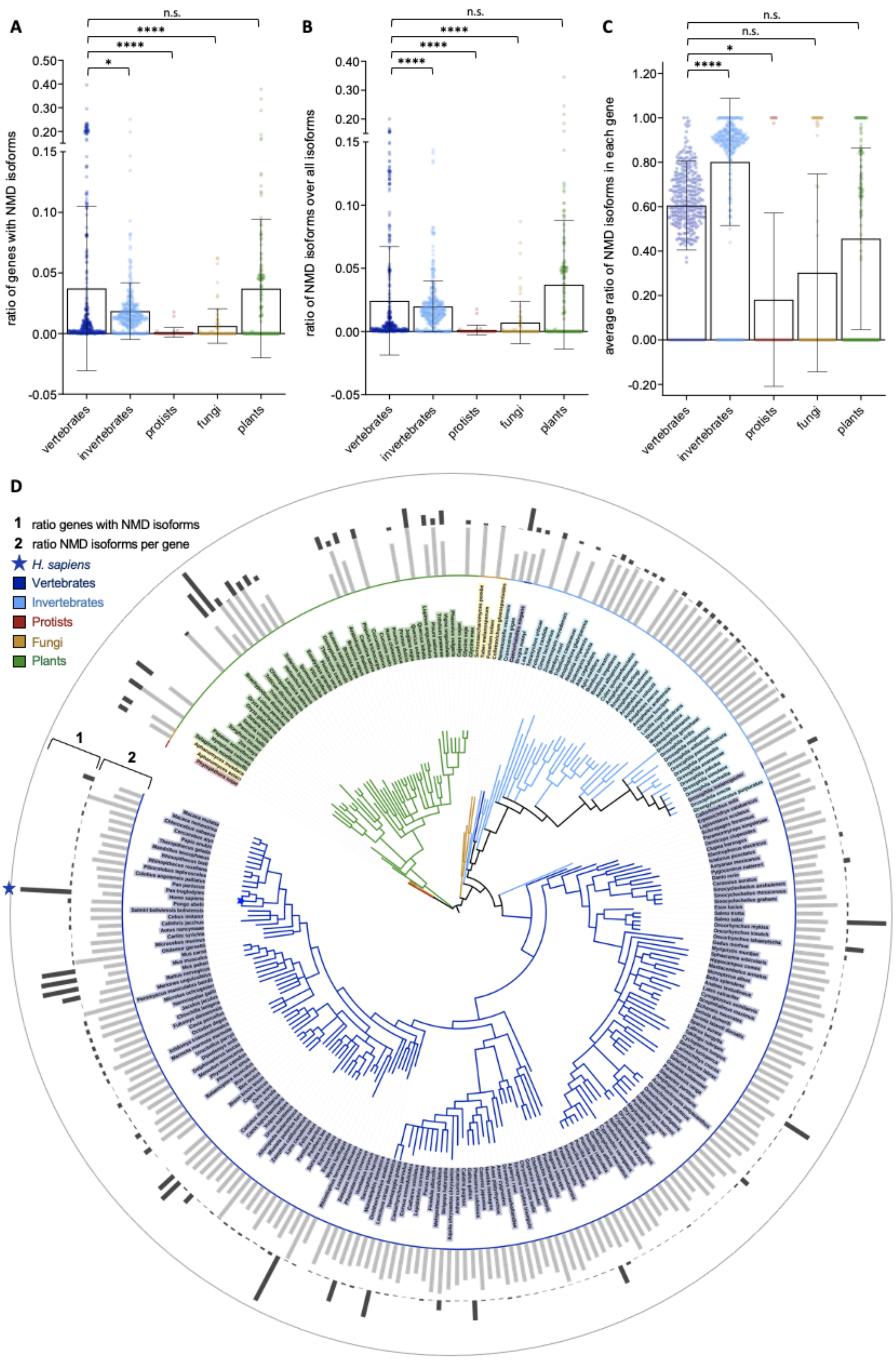


**Supplementary Figure S1.** Analysis of annotated NMD isoforms across different taxonomic groups (vertebrates, invertebrates, protists, fungi and plants). (**A**) Ratio of coding genes with at least one annotated NMD isoform for each species. (**B**) Ratio of all annotated NMD isoforms over all annotated isoforms for each species. (**C**) Average ratio calculated for each species between annotated NMD isoforms and all annotated isoforms of each coding gene. In all plots, each dot represents a species, and the median is indicated as a black horizontal line. Statistical significance by non-parametric Kruskal–Wallis test followed by Dunn's multiple comparisons test is shown for the vertebrate group compared to all others. * $p<0.05$, ** $p<0.01$, *** $p<0.001$, **** $p<0.0001$, ns: not significant. (**D**) NMD isoforms annotated throughout the genomes of eukaryotic species that were filtered for a BUSCO completeness score of the annotation ≥ 90%. For each species, bars indicate either the ratio of coding genes with at least one annotated NMD isoform (Track "1", highest value: 0.38), or the average ratio of annotated NMD isoforms over all isoforms of each coding gene (Track "2", highest value: 0.57).

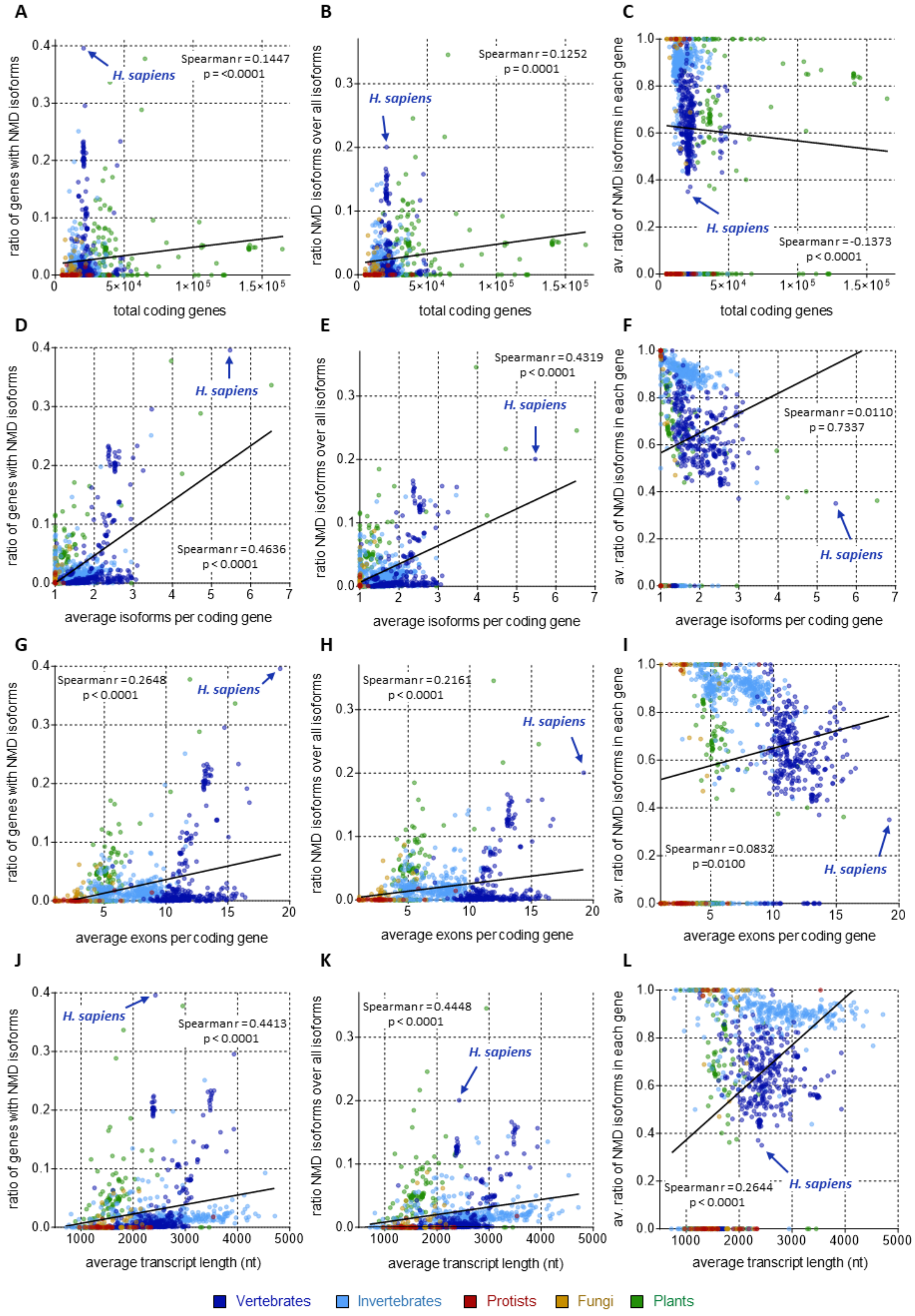
A
ratio of genes with NMD isoforms
H. sapiens
Spearman r = 0.1447
p = <0.0001
total coding genes
B
ratio NMD isoforms over all isoforms
H. sapiens
Spearman r = 0.1252
p = 0.0001
total coding genes
C
av. ratio of NMD isoforms in each gene
H. sapiens
Spearman r = -0.1373
p < 0.0001
total coding genes
D
ratio of genes with NMD isoforms
H. sapiens
Spearman r = 0.4636
p < 0.0001
average isoforms per coding gene
E
ratio NMD isoforms over all isoforms
Spearman r = 0.4319
p < 0.0001
H. sapiens
average isoforms per coding gene
F
av. ratio of NMD isoforms in each gene
Spearman r = 0.0110
p = 0.7337
H. sapiens
average isoforms per coding gene
G
ratio of genes with NMD isoforms
Spearman r = 0.2648
p < 0.0001
H. sapiens
average exons per coding gene
H
ratio NMD isoforms over all isoforms
Spearman r = 0.2161
p < 0.0001
H. sapiens
average exons per coding gene
I
av. ratio of NMD isoforms in each gene
Spearman r = 0.0832
p = 0.0100
H. sapiens
average exons per coding gene
J
ratio of genes with NMD isoforms
H. sapiens
Spearman r = 0.4413
p < 0.0001
average transcript length (nt)
K
ratio NMD isoforms over all isoforms
Spearman r = 0.4448
p < 0.0001
H. sapiens
average transcript length (nt)
L
av. ratio of NMD isoforms in each gene
Spearman r = 0.2644
p < 0.0001
H. sapiens
average transcript length (nt)
Vertebrates
Invertebrates
Protists
Fungi
Plants

36

**Supplementary Figure S2.** The ratio of coding genes with at least one annotated NMD isoform, the ratio of all annotated NMD isoforms over all annotated isoforms, and the average ratio between annotated NMD isoforms and all annotated isoforms of each coding gene are respectively compared for each species with (**A**, **B** and **C**) the total number of annotated coding genes, (**D**, **E** and **F**) the average number of isoforms annotated for each coding gene, (**G**, **H** and **I**) the average number of exons annotated for each coding gene, and (**J**, **K** and **L**) the average length (nt) of annotated coding transcripts. In all plots, each dot represents a species. Simple linear regression is shown for each scatter plot as a black line, with relative Spearman correlation coefficient (r) and p-value (p).

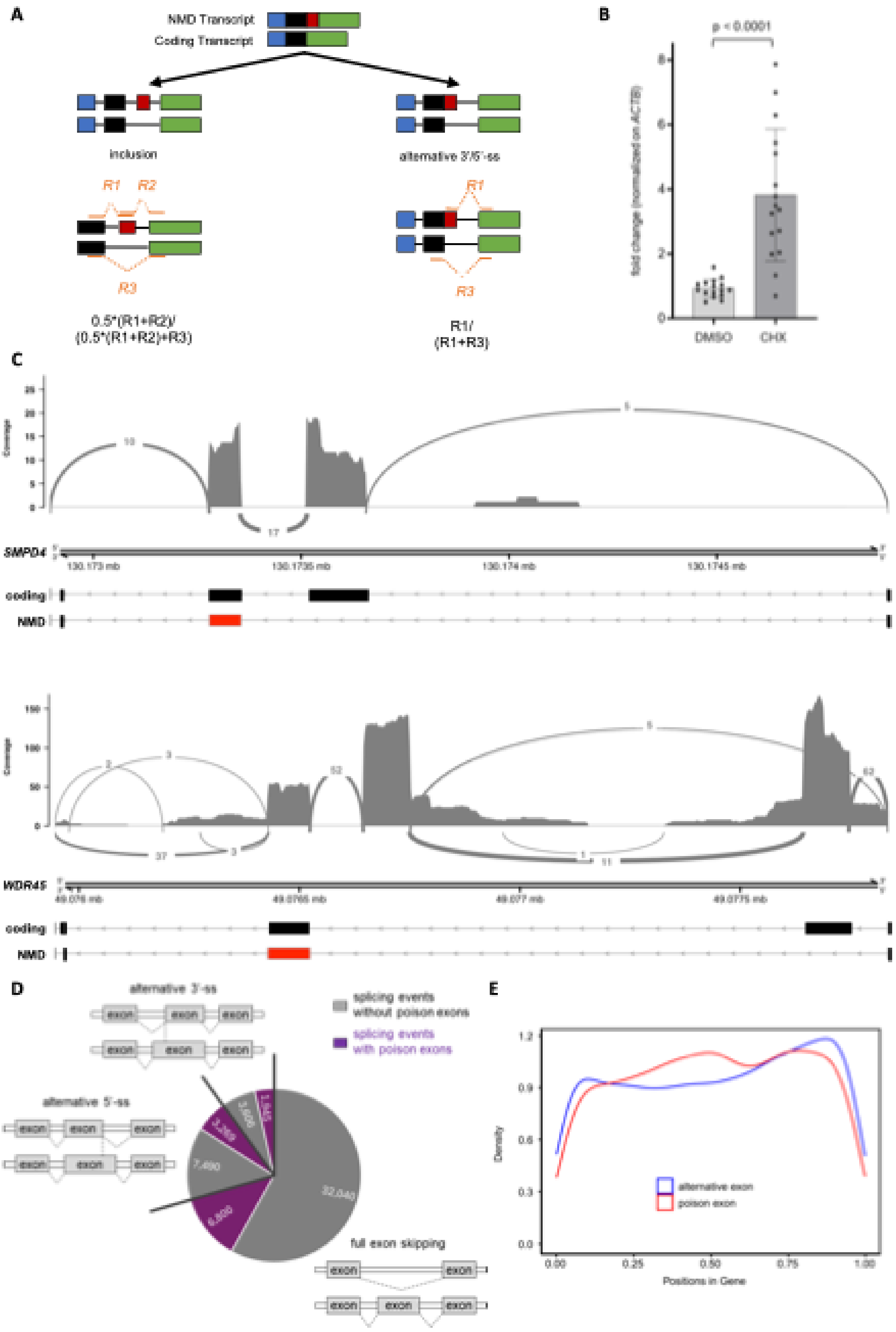
A
NMD Transcript
Coding Transcript
inclusion
alternative 3'/5'-ss
R1
R2
R3
0.5*(R1+R2)/
(0.5*(R1+R2)+R3)
R1/
(R1+R3)
B
p < 0.0001
fold change (normalized on ACTB)
DMSO
CHX
C
Coverage
SMPD4
coding
NMD
WDR45
D
alternative 3'-ss
alternative 5'-ss
full exon skipping
exon
splicing events without poison exons
splicing events with poison exons
32,040
7,490
6,800
3,269
3,606
1,842
E
Density
Positions in Gene
alternative exon
poison exon

51

**Supplementary Figure S3.** (**A**) Schematic representation of different classes of alternatively spliced PEs (“inclusion”, and “alternative 3’/5’-ss”) and PSI calculation methods. PEs are shown as red boxes. R1, R2, and R3 represent reads aligned to a given transcript region. (**B**) Sensitivity to NMD is assessed by CHX treatment in HEK293T cells for 16 different transcripts carrying putative PEs. Each couple of connected dots represents a different transcript. Cells were treated with 50 µg/mL CHX (or DMSO as control), and the increase of transcript levels was evaluated by RT-qPCR. Primers specific for the exon-exon junction that includes the PE were used for each transcript. All values are internally normalized on the expression of *ACTB* gene, while transcript levels are represented as fold change compared to the DMSO condition. Statistical significance by Student’s t-test is shown for the comparison between the DMSO and CHX conditions. Data are shown as average ± standard deviation. Each data point represents a different gene. (C) Sashimi plots for PEs of genes *SMPD4* and *WDR45* from HEK293T cells. For each panel, the top track shows the read coverage and splice junction reads from RNA-seq, with arc thickness representing the number of supporting split reads. The middle track shows the genomic coordinates of the locus. The bottom tracks show the relevant transcript isoforms: the protein-coding transcript and the PE-containing transcript that is targeted by NMD. In all panels, red boxes denote the poison exon (PE) and black boxes denote flanking exons, and arrows indicate the direction of transcription. (**D**) Different types of alternative splicing events with or without putative PEs are counted across the human genome annotation. They include canonical exon exclusion/inclusion events and alternative 3’- and 5’-SSs. (**E**) Relative positions of putative PEs and canonical alternative exons occurrence along the host genes, calculated as the distance from TSS divided by the gene length.

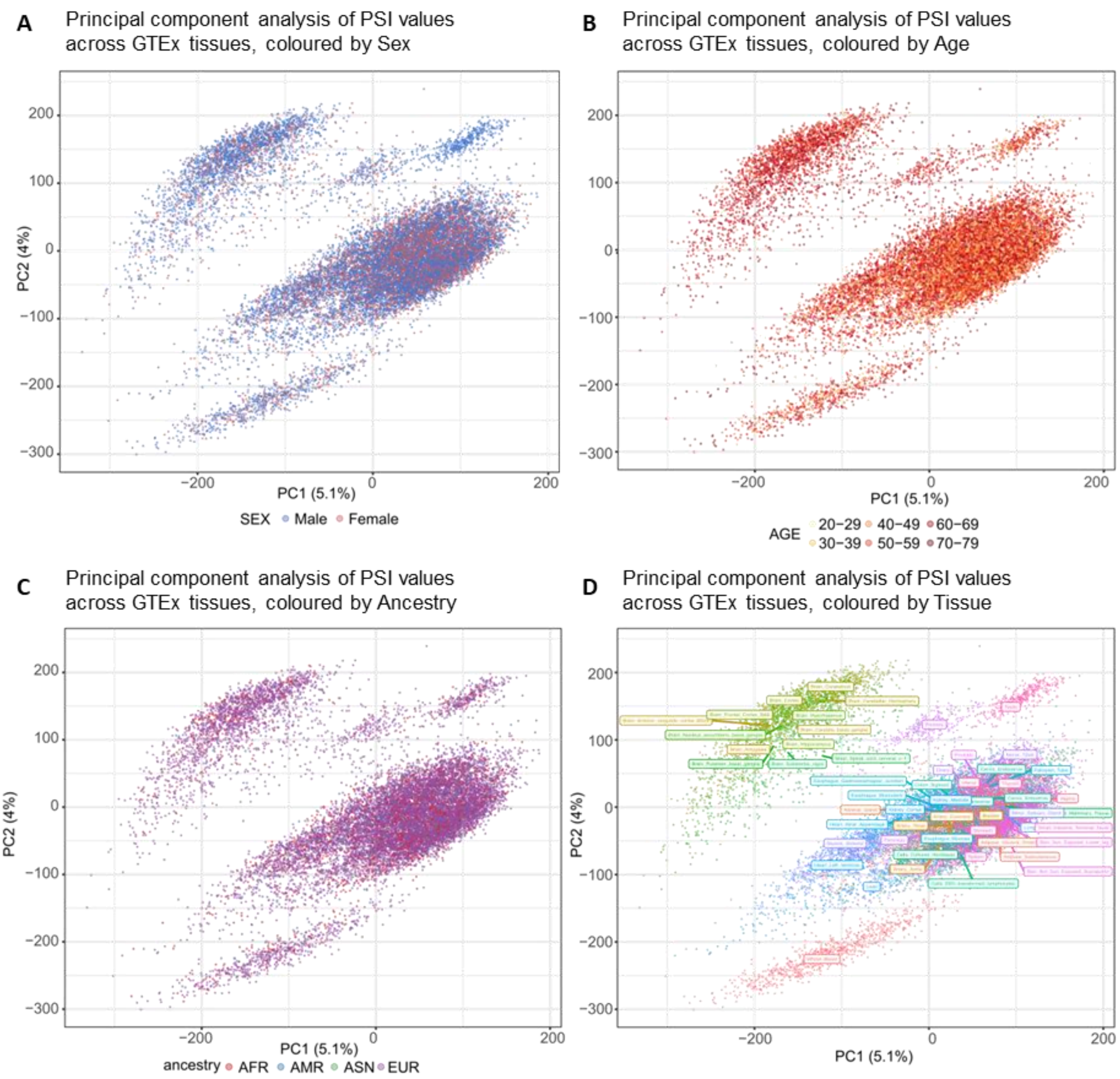


**Supplementary Figure S4.** Principal component analysis (PCA) plot of demographic covariates on PSI matrix across GTEx tissues. Samples are colored by **(A)** sex, **(B)** age group, **(C)** ancestry, and **(D)** tissue type.

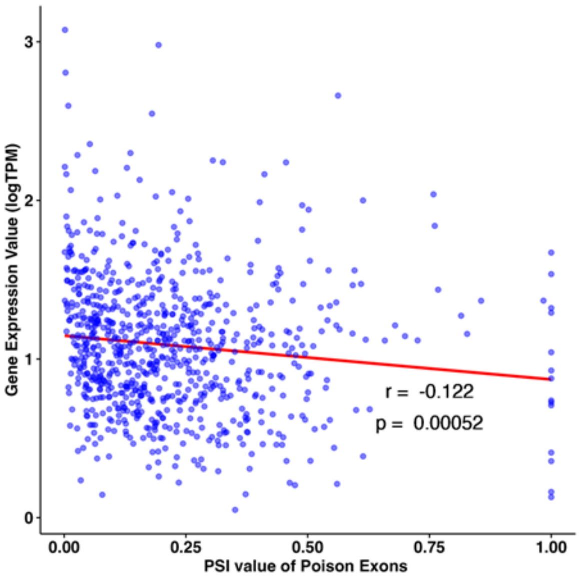


**Supplementary Figure S5**. Correlation between PE's inclusion (as PSI value, x-axis) and its host gene's expression (as TPM value, y-axis) in the human brain based on GTEx data. Each dot represents a PE. Simple linear regression is shown as a red line, with relative Pearson correlation coefficient (r) and p-value (p).

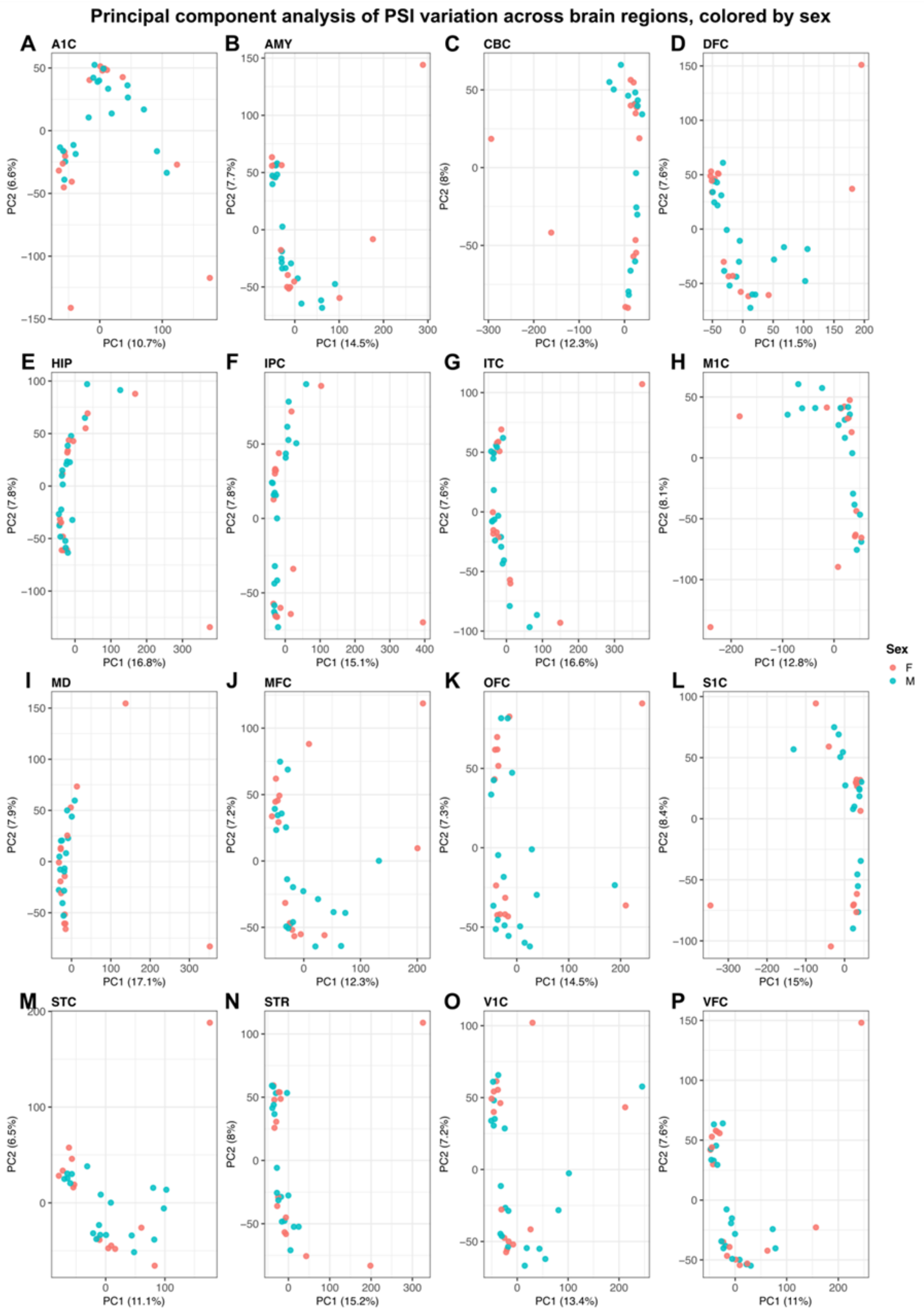


**Supplementary Figure S6**. Principal component analysis of PE inclusion across BrainSpan samples, colored by Sex. (**A-P**) Each panel shows PC1 versus PC2 for one brain region, with samples colored by Sex. The percent variance explained by each principal component is indicated in the axis labels. A1C: Primary auditory cortex; AMY: Amygdala; CBC: Cerebellar cortex; DFC: Dorsolateral prefrontal cortex; HIP: Hippocampus; IPC: Posterior inferior parietal cortex; ITC: inferior temporal cortex; MD: Mediodorsal nucleus of the thalamus; MFC: Medial prefrontal cortex; OFC: Orbital prefrontal cortex; S1C: Primary somatosensory cortex; STR: Striatum; V1C: Primary visual cortex; M1C: Primary motor cortex; VFC: Ventrolateral prefrontal cortex. STC: Superior temporal cortex.

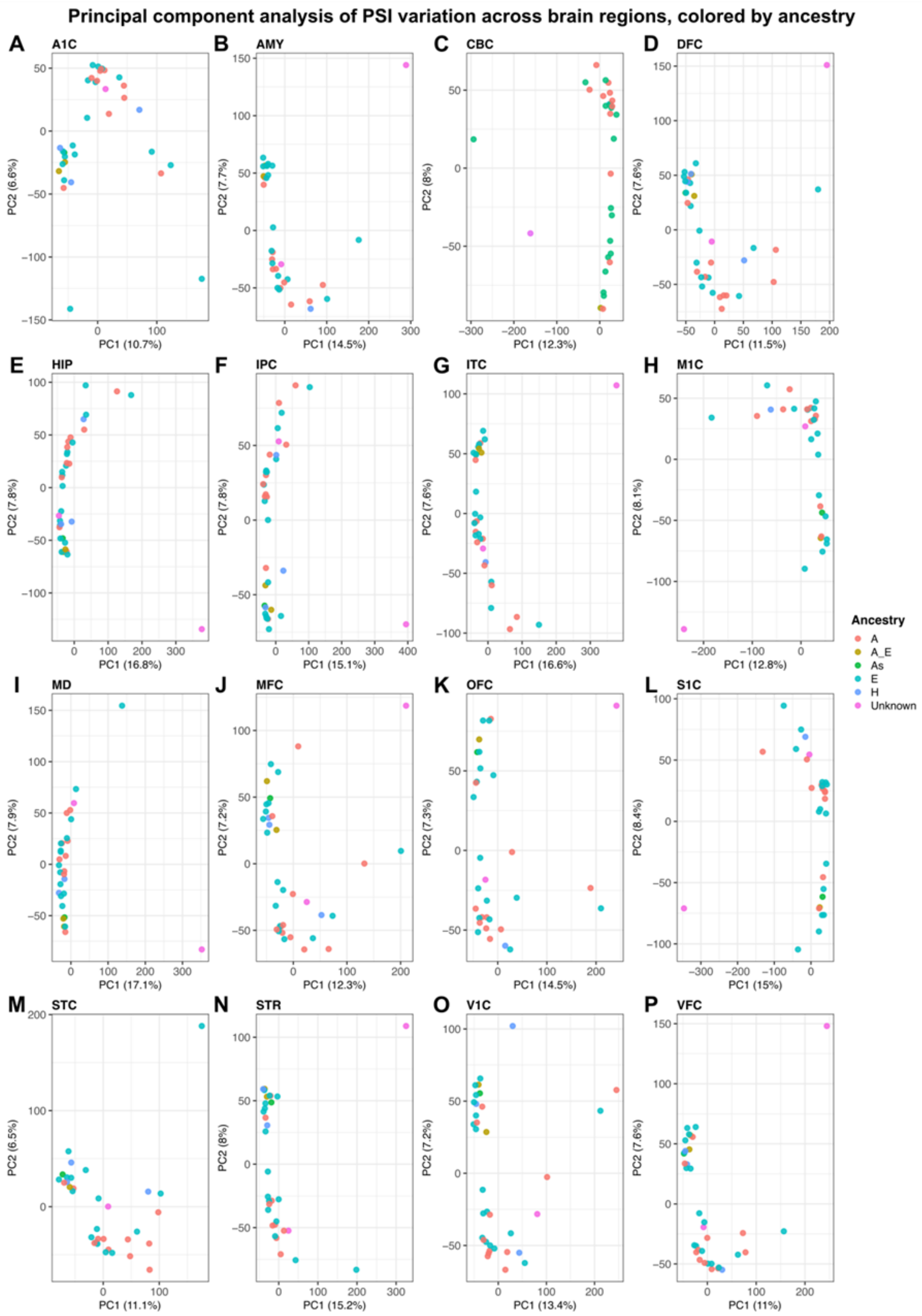


**Supplementary Figure S7**. Principal component analysis of PE inclusion across BrainSpan samples, colored by ancestry. (**A-P**) Each panel shows PC1 versus PC2 for one brain region, with samples colored by ancestry. The percent variance explained by each principal component is indicated in the axis labels. A1C: Primary auditory cortex; AMY: Amygdala; CBC: Cerebellar cortex; DFC: Dorsolateral prefrontal cortex; HIP: Hippocampus; IPC: Posterior inferior parietal cortex; ITC: inferior temporal cortex; MD: Mediodorsal nucleus of the thalamus; MFC: Medial prefrontal cortex; OFC: Orbital prefrontal cortex; S1C: Primary somatosensory cortex; STR: Striatum; V1C: Primary visual cortex; M1C: Primary motor cortex; VFC: Ventrolateral prefrontal cortex. STC: Superior temporal cortex.

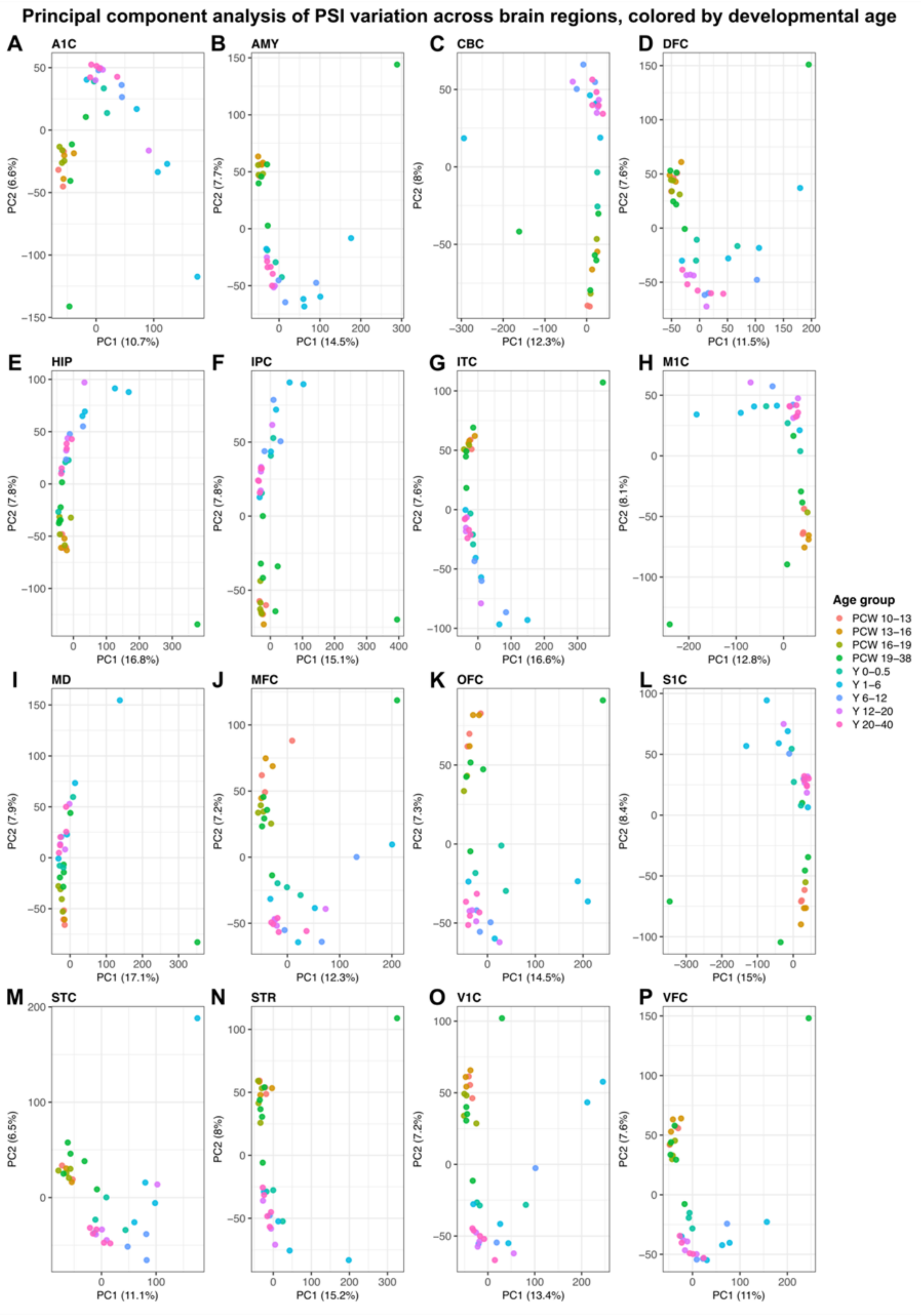


**Supplementary Figure S8**. Principal component analysis of PE inclusion across BrainSpan samples, colored by developmental age. (**A-P**) Each panel shows PC1 versus PC2 for one brain region, with samples colored by developmental age group. The percent variance explained by each principal component is indicated in the axis labels. A1C: Primary auditory cortex; AMY: Amygdala; CBC: Cerebellar cortex; DFC: Dorsolateral prefrontal cortex; HIP: Hippocampus; IPC: Posterior inferior parietal cortex; ITC: inferior temporal cortex; MD: Mediodorsal nucleus of the thalamus; MFC: Medial prefrontal cortex; OFC: Orbital prefrontal cortex; S1C: Primary somatosensory cortex; STR: Striatum; V1C: Primary visual cortex; M1C: Primary motor cortex; VFC: Ventrolateral prefrontal cortex. STC: Superior temporal cortex.

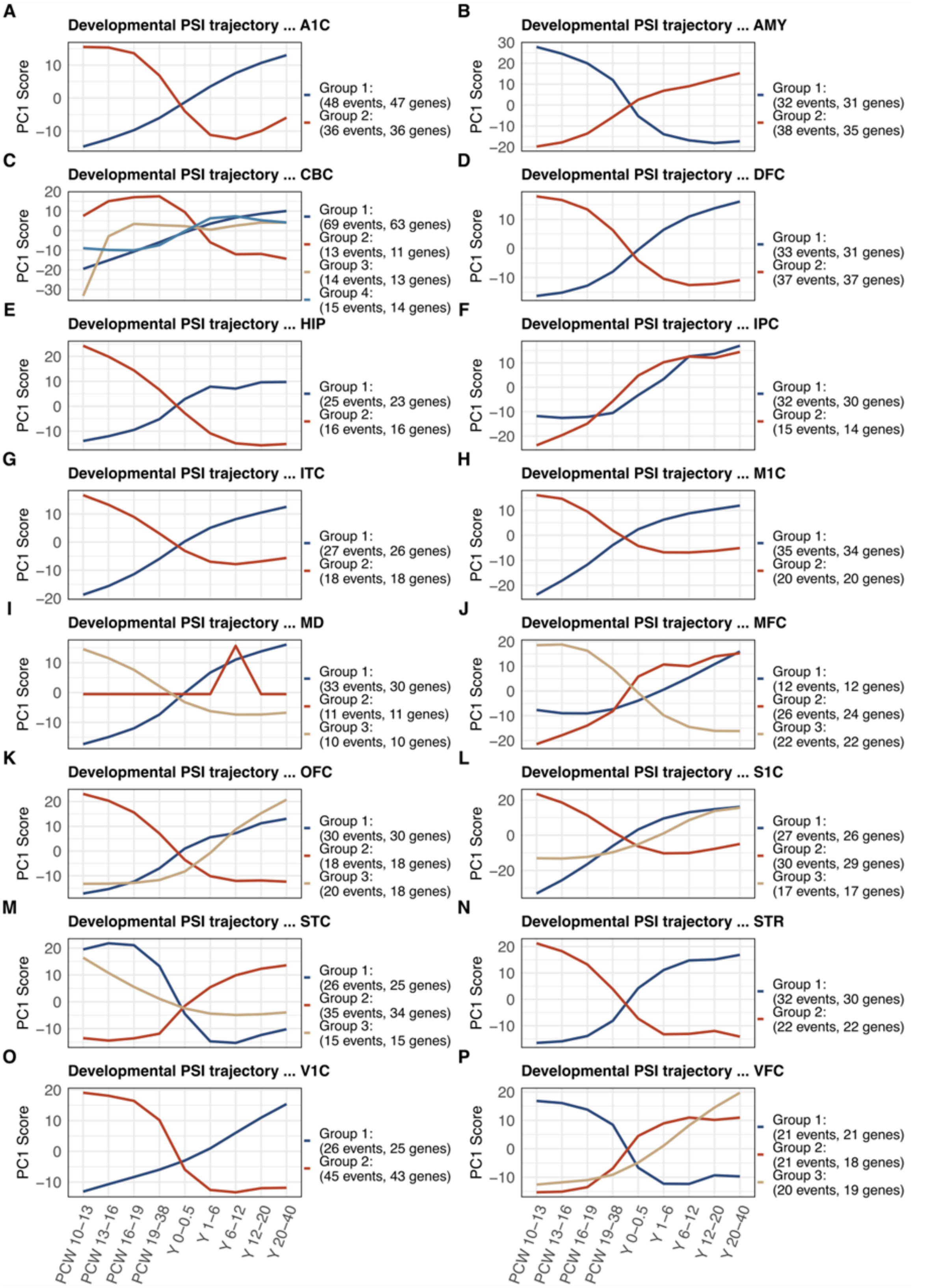


**Supplementary Figure S9**. (**A-P**) PE inclusion clustered into trajectories in different brain regions. The two numbers in the parentheses represent total PE events and host genes, respectively, in each cluster. A1C: Primary auditory cortex; AMY: Amygdala; CBC: Cerebellar cortex; DFC: Dorsolateral prefrontal cortex; HIP: Hippocampus; IPC: Posterior inferior parietal cortex; ITC: inferior temporal cortex; MD: Mediodorsal nucleus of the thalamus; MFC: Medial prefrontal cortex; OFC: Orbital prefrontal cortex; S1C: Primary somatosensory cortex; STR: Striatum; V1C: Primary visual cortex; M1C: Primary motor cortex; VFC: Ventrolateral prefrontal cortex. STC: Superior temporal cortex.

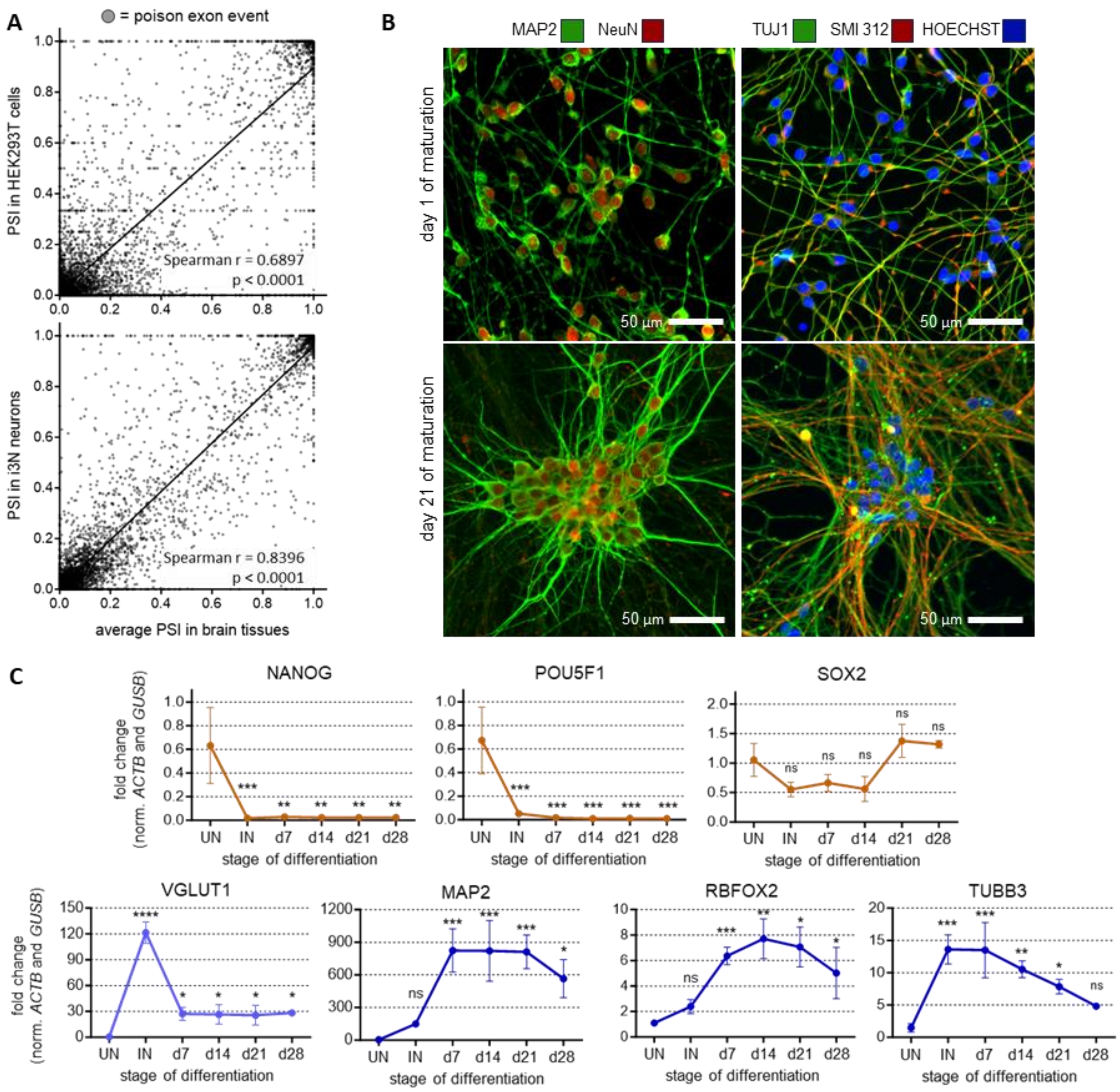


**Supplementary Figure S10**. (**A**) PSI values of putative PEs are correlated between the human brain and HEK293T cells (top) or i3N neurons (bottom). Simple linear regression is shown as a black line, with relative Spearman correlation coefficient (r) and p-value (p). (**B**) Expression of pluripotency (orange lines) and neuronal differentiation (blue lines) markers is assessed by RT-qPCR at the stage of undifferentiated iPSCs ("UN"), induced neurons following 3 days of NGN2 overexpression ("IN"), and neurons at 7, 14, 21 or 28 days of maturation. All values are internally normalized on the expression of genes *ACTB* and *GUSB*, while expression values are represented as fold-change compared to the "UN" condition. Statistical significance is indicated for each stage compared to "UN". * $p<0.05$, ** $p<0.01$, *** $p<0.001$, **** $p<0.0001$, ns: not significant. Data are shown as average of three replicates ± standard deviation. (**C**) Immunocytochemistry analysis with antibodies for neuronal markers MAP2, RBFOX3 (NeuN), TUBB3 (TUJ1), and neurofilaments (SMI 312) using neurons at 1 or 21 days of maturation. Images were taken with 20x magnification.

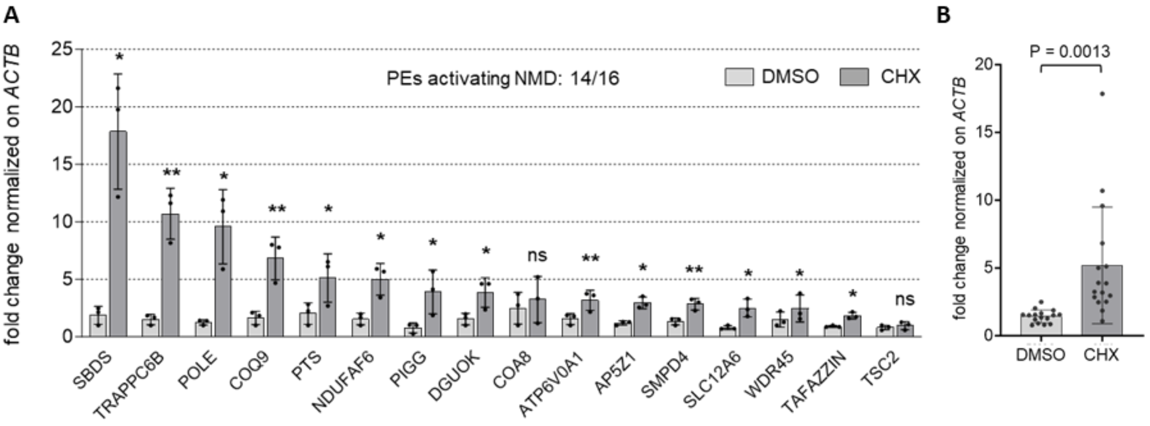


**Supplementary Figure S11**. (**A** and **B**) Sensitivity to NMD is assessed by CHX treatment in i3N neurons for 16 different transcripts carrying putative PEs. The analysis is shown for every single transcript (figure A) and for all transcripts together (with couple of connected dots representing a different transcript, figure B). Neurons were treated with 50 µg/mL CHX (or DMSO as control) at 21 days of maturation, and the increase of transcript levels was evaluated by RT-qPCR. Primers specific for the exon-exon junction that includes the PE were used for each transcript. All values are internally normalized on the expression of genes *ACTB* and *GUSB*, while transcript levels are represented as fold change compared to the DMSO condition. Statistical significance by Student's t-test is indicated for each comparison. Data are shown as average ± standard deviation. Each data point represents one of three replicates (A) or a different gene (B).

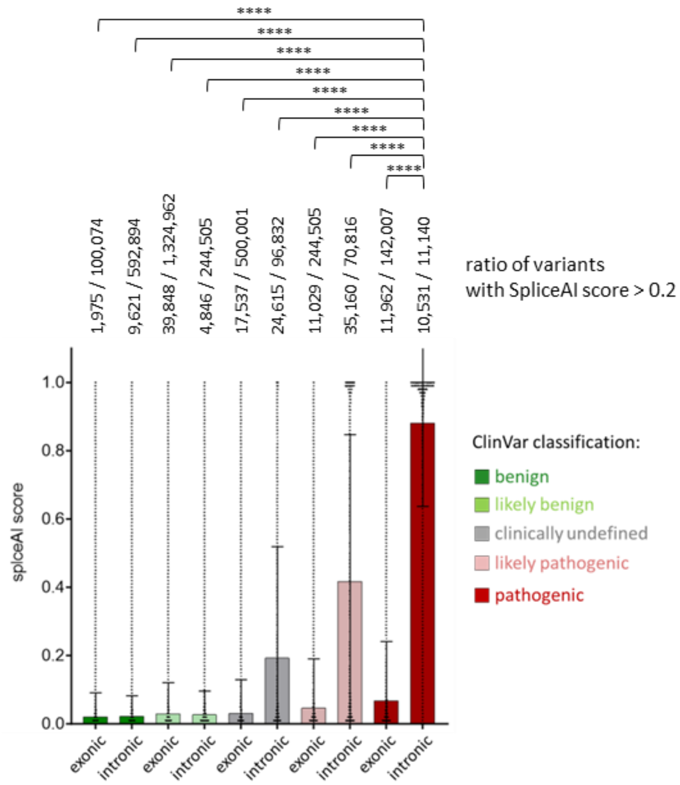


**Supplementary Figure S12**. SpliceAI prediction of the impact of 236,304 ClinVar genetic variants on the nearby splice sites. Variants are grouped based on their annotated pathogenicity ("benign", "likely benign", "undefined", "pathogenic", and "likely pathogenic") and their location within exonic or intronic regions. The ratio of variants with SpliceAI score > 0.2 is shown at the top of each bar. Statistical significance by non-parametric Kruskal–Wallis test, followed by Dunn's multiple comparisons test, is shown for the "intronic pathogenic" group compared to all others. * p<0.05, ** p<0.01, *** p<0.001, **** p<0.0001, ns: not significant.

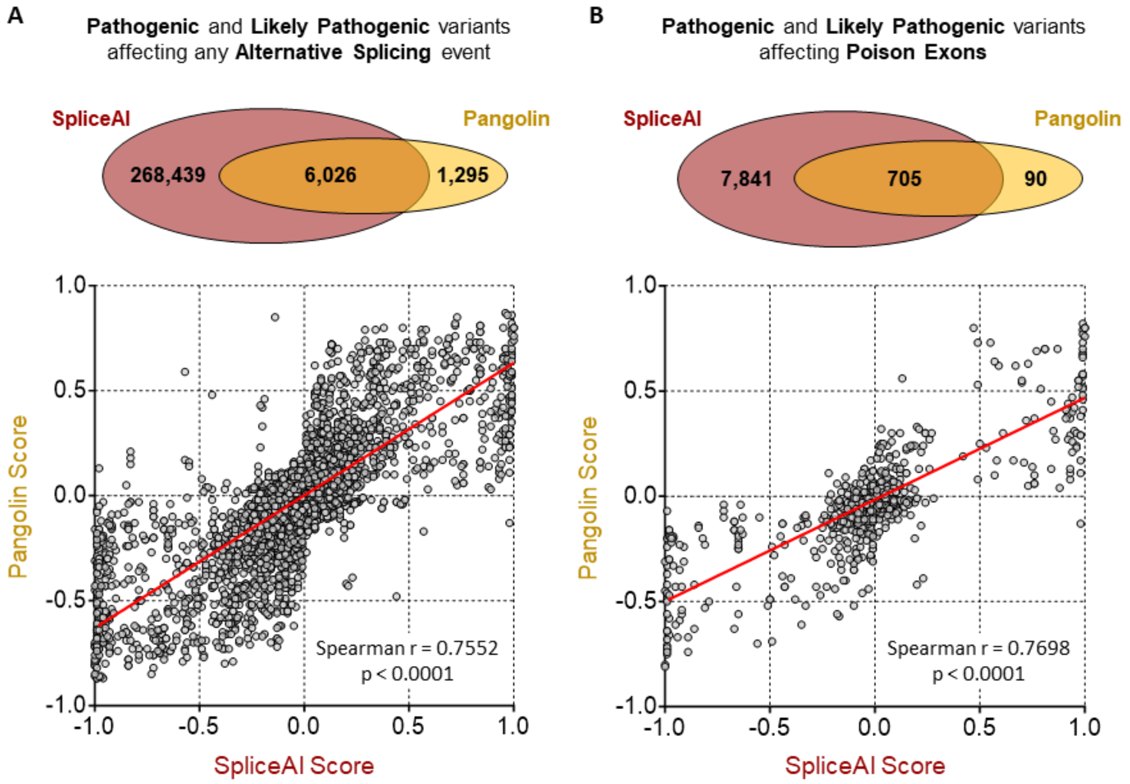


**Supplementary Figure S13.** The effect on (**A**) any alternative splicing event or (**B**) PE-including splicing events is predicted for pathogenic and likely-pathogenic mutations from Clinvar using either SpiceAI or Pangolin. Venn diagrams at the top show predicted splicing mutations common or unique to SpliceAI and Pangolin. Scatter plots at the bottom show the correlation between SpliceAI score and Pangolin score for common splicing mutations. Simple linear regression is shown for each scatter plot as a black line, with relative Spearman correlation coefficient (r) and p-value (p).

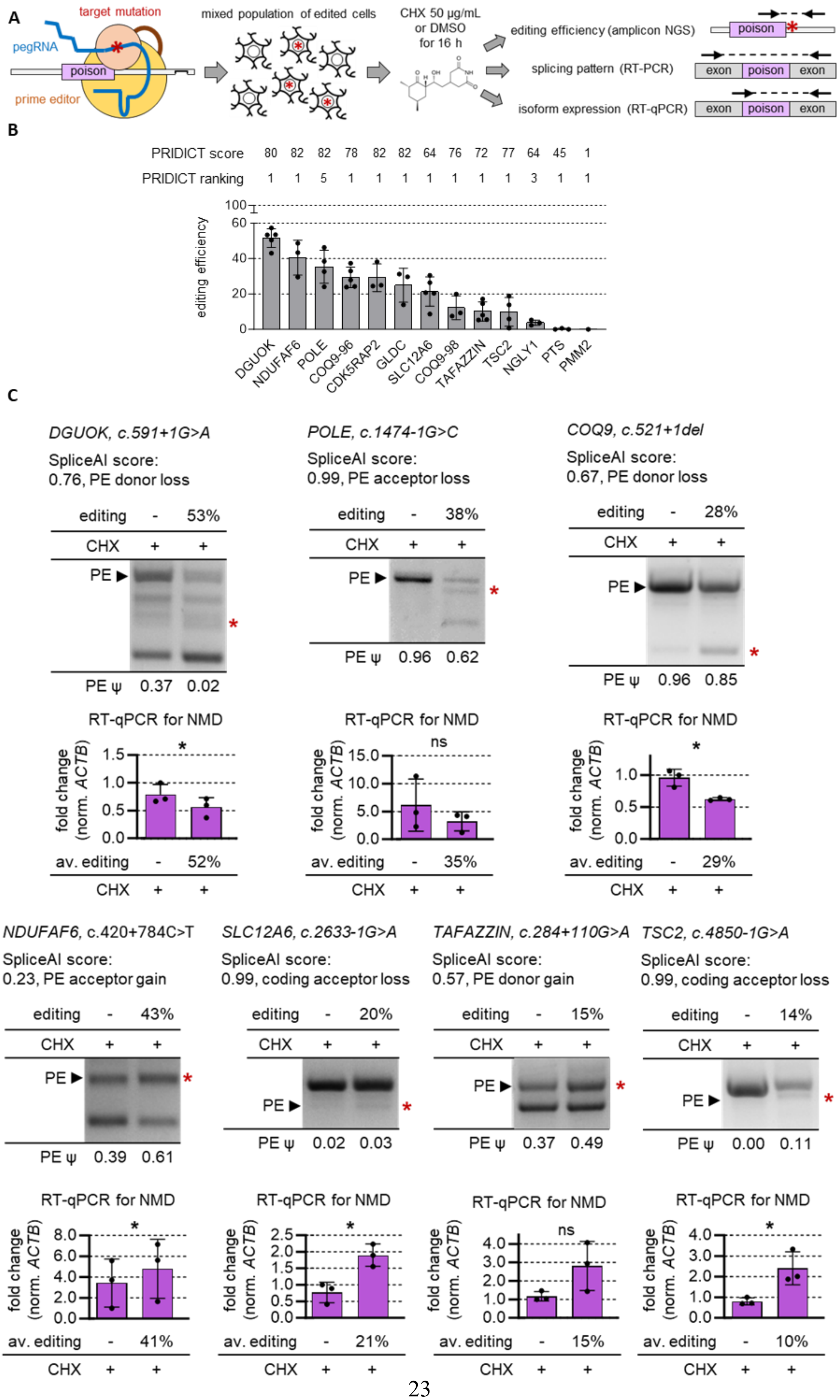
A
target mutation
pegRNA
poison
prime editor
mixed population of edited cells
CHX 50 µg/mL
or DMSO
for 16 h
editing efficiency (amplicon NGS)
splicing pattern (RT-PCR)
isoform expression (RT-qPCR)
exon
poison
exon
B
PRIDICT score 80 82 82 78 82 82 64 76 72 77 64 45 1
PRIDICT ranking 1 1 5 1 1 1 1 1 1 1 3 1 1
editing efficiency
DGUOK
NDUFAF6
POLE
COQ9-96
CDK5RAP2
GLDC
SLC12A6
COQ9-98
TAFAZZIN
TSC2
NGLY1
PTS
PMM2
C
DGUOK, c.591+1G>A
SpliceAI score:
0.76, PE donor loss
editing - 53%
CHX + +
PE ψ 0.37 0.02
RT-qPCR for NMD
fold change (norm. ACTB)
av. editing - 52%
CHX + +
POLE, c.1474-1G>C
SpliceAI score:
0.99, PE acceptor loss
editing - 38%
CHX + +
PE ψ 0.96 0.62
RT-qPCR for NMD
av. editing - 35%
CHX + +
COQ9, c.521+1del
SpliceAI score:
0.67, PE donor loss
editing - 28%
CHX + +
PE ψ 0.96 0.85
RT-qPCR for NMD
av. editing - 29%
CHX + +
NDUFAF6, c.420+784C>T
SpliceAI score:
0.23, PE acceptor gain
editing - 43%
CHX + +
PE ψ 0.39 0.61
RT-qPCR for NMD
av. editing - 41%
CHX + +
SLC12A6, c.2633-1G>A
SpliceAI score:
0.99, coding acceptor loss
editing - 20%
CHX + +
PE ψ 0.02 0.03
RT-qPCR for NMD
av. editing - 21%
CHX + +
TAFAZZIN, c.284+110G>A
SpliceAI score:
0.57, PE donor gain
editing - 15%
CHX + +
PE ψ 0.37 0.49
RT-qPCR for NMD
av. editing - 15%
CHX + +
TSC2, c.4850-1G>A
SpliceAI score:
0.99, coding acceptor loss
editing - 14%
CHX + +
PE ψ 0.00 0.11
RT-qPCR for NMD
av. editing - 10%
CHX + +

**Supplementary Figure S14**. (**A**) Representation of the experimental workflow used to validate the effect of genetic variants on PE splicing. (**B**) Editing efficiencies of 13 different pegRNAs, measured by amplicon-NGS in HEK293T cells. (**C**) Splicing pattern analysis by agarose-gel RT-PCR (top panels) and RT-qPCR (bottom panels) is shown for 7 genetic variants that have been successfully introduced into HEK293T cells by prime editing. Cells were treated with CHX 16 h prior to collection. The percentage of edited cells, measured by amplicon-NGS, is reported for each experiment. Bands corresponding to the PE-containing RNA isoform are indicated by a black arrowhead. Bands corresponding to isoforms predicted to increase upon mutation are indicated with an asterisk. PSI of the PE-containing isoform (calculated by densitometry analysis) is reported for each lane. Primers specific for the NMD isoform were used for the RT-qPCR. All values are internally normalized on the expression of *ACTB* gene, while transcript levels are represented as fold change compared to the DMSO condition with no editing. The percentage of edited cells, measured by amplicon-NGS, is reported for each experiment as the average of multiple replicates. Statistical significance by Student's t-test is shown for each comparison. * $p<0.05$, ** $p<0.01$, *** $p<0.001$, **** $p<0.0001$, ns: not significant. Data are shown as average ± standard deviation. Each data point represents one of three replicates.

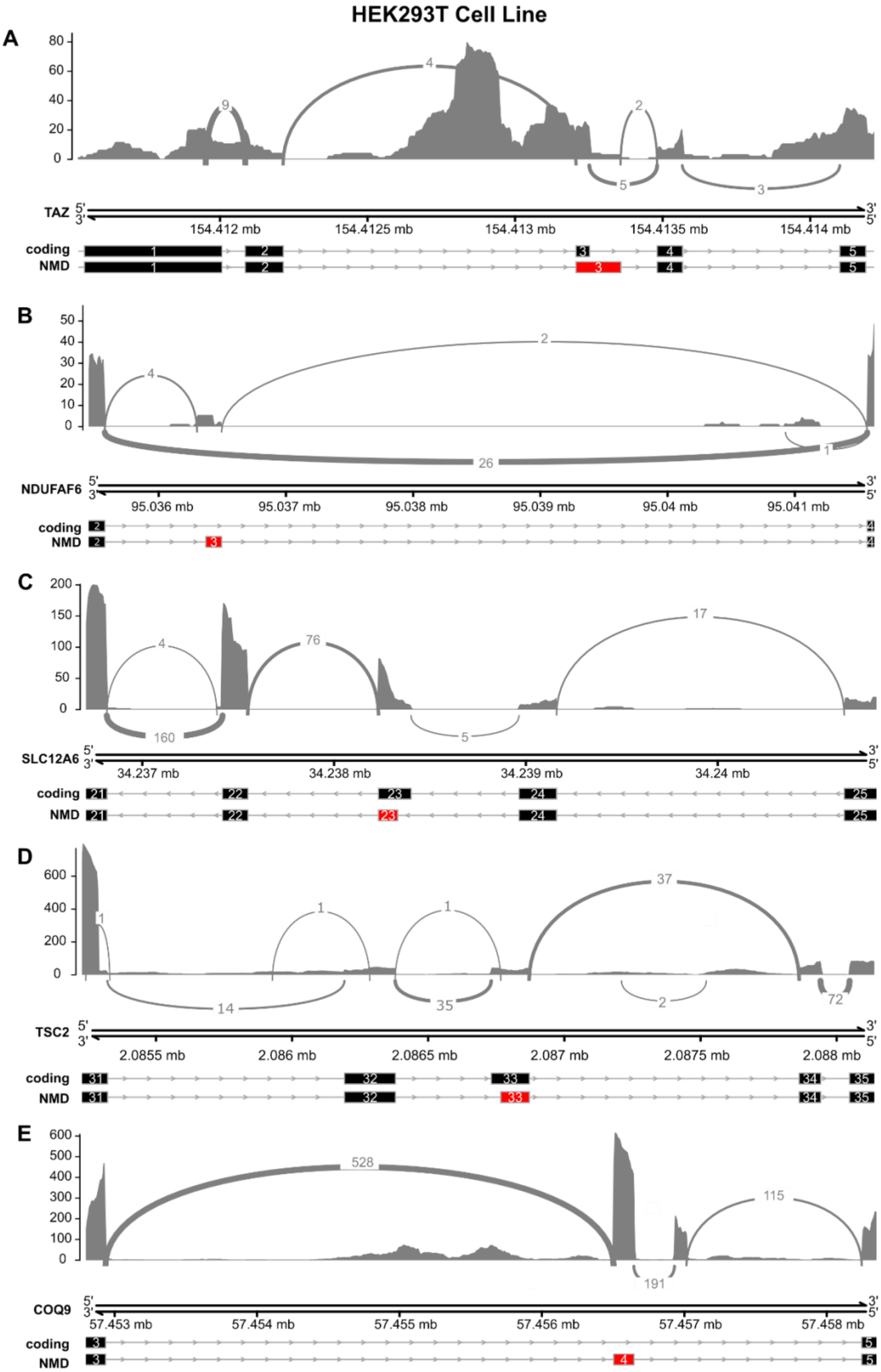
HEK293T Cell Line
A
TAZ
154.412 mb
154.4125 mb
154.413 mb
154.4135 mb
154.414 mb
coding
NMD
B
NDUFAF6
95.036 mb
95.037 mb
95.038 mb
95.039 mb
95.04 mb
95.041 mb
coding
NMD
C
SLC12A6
34.237 mb
34.238 mb
34.239 mb
34.24 mb
coding
NMD
D
TSC2
2.0855 mb
2.086 mb
2.0865 mb
2.087 mb
2.0875 mb
2.088 mb
coding
NMD
E
COQ9
57.453 mb
57.454 mb
57.455 mb
57.456 mb
57.457 mb
57.458 mb
coding
NMD

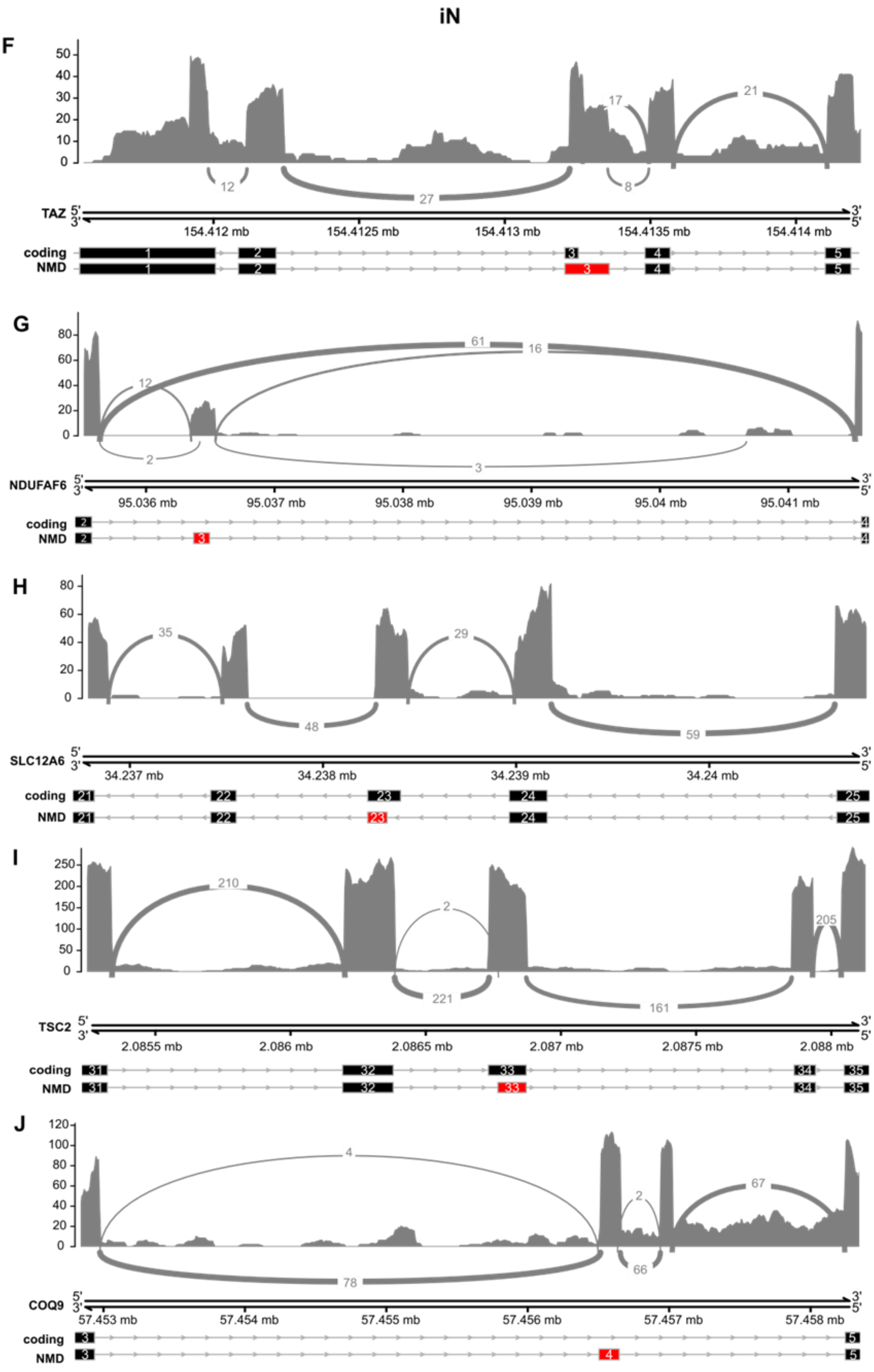
iN
F
TAZ
coding
NMD
154.412 mb
154.4125 mb
154.413 mb
154.4135 mb
154.414 mb
G
NDUFAF6
coding
NMD
95.036 mb
95.037 mb
95.038 mb
95.039 mb
95.04 mb
95.041 mb
H
SLC12A6
coding
NMD
34.237 mb
34.238 mb
34.239 mb
34.24 mb
I
TSC2
coding
NMD
2.0855 mb
2.086 mb
2.0865 mb
2.087 mb
2.0875 mb
2.088 mb
J
COQ9
coding
NMD
57.453 mb
57.454 mb
57.455 mb
57.456 mb
57.457 mb
57.458 mb

**Supplementary Figure S15**. Sashimi plots for selected PEs of genes *TAZ*, *NDUFAF6*, *SLC12A6*, *TARDBP*, *TSC2* and *COQ9*. **(A–E)** HEK293T cells: sashimi plots for TAZ (A), NDUFAF6 (B), SLC12A6 (C), TSC2 (D), and COQ9 (E). (F–J) iPSC-derived neurons (iNs): corresponding plots for the same genes in the same order. For each panel, the top track shows the read coverage and splice junction reads from RNA-seq, with arc thickness representing the number of supporting split reads. The middle track shows the genomic coordinates of the locus. The bottom tracks show the relevant transcript isoforms: the protein-coding transcript and the PE-containing transcript that is targeted by NMD. In all panels, red boxes denote the poison exon (PE) and black boxes denote flanking exons, and arrows indicate the direction of transcription.

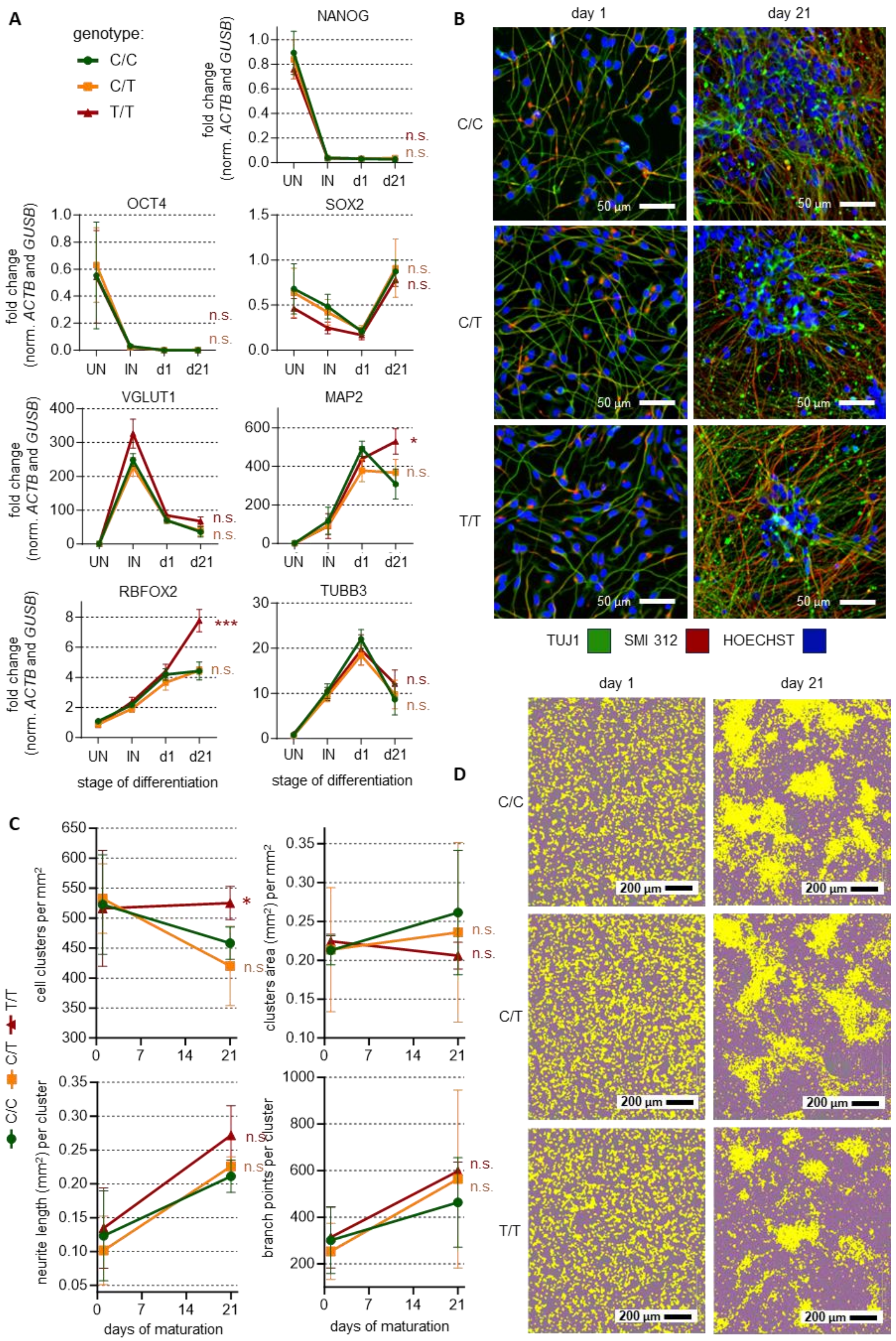
A
genotype:
C/C
C/T
T/T
NANOG
OCT4
SOX2
VGLUT1
MAP2
RBFOX2
TUBB3
fold change
(norm. ACTB and GUSB)
UN
IN
d1
d21
n.s.
*
***
stage of differentiation
B
day 1
day 21
C/C
C/T
T/T
50 µm
TUJ1
SMI 312
HOECHST
C
cell clusters per mm²
clusters area (mm²) per mm²
neurite length (mm²) per cluster
branch points per cluster
days of maturation
D
day 1
day 21
200 µm

**Supplementary Figure S16**. (**A**) Expression of pluripotency and neuronal differentiation markers is assessed by RT-qPCR for c.420+784C>T clonal cell lines at the stage of undifferentiated iPSCs ("UN"), induced neurons following 3 days of NGN2 overexpression ("IN"), and neurons at 1 or 21 days of maturation. All values are internally normalized on the expression of genes *ACTB* and *GUSB*, while expression values are represented as fold-change compared to the "UN" condition of the wild-type line. Statistical significance by one-way ANOVA with Geisser-Greenhouse correction is indicated for the d21 stage of the mutant lines compared to wild-type. * $p<0.05$, ** $p<0.01$, *** $p<0.001$, **** $p<0.0001$, ns: not significant. Data are shown as average of three replicates ± standard deviation. (**B**) Immunocytochemistry analysis of c.420+784C>T clonal cell lines, at 1 or 21 days of maturation, using antibodies for neuronal markers TUBB3 (TUJ1) and neurofilaments (SMI 312). Hoechst is used for nuclear staining. (**C**) Quantitative analysis of the number (per $mm^2$) and size (ratio of covered area) of cell-body clusters, the length of neurites (normalized with the number of clusters), and the number of neurite branch points (per cell cluster) for the c.420+784C>T clonal cell cultures at 1 or 21 days of neuronal maturation. Statistical significance by one-way ANOVA with Geisser-Greenhouse correction is shown for each comparison. Data are shown as average of three replicates ± standard deviation. (**D**) Neurites (purple) and cell bodies (yellow) are highlighted for clonal cell cultures with different genotypes for c.420+784C>T at 1 or 21 days of neuronal maturation. Images are representative of multiple fields and replicates.
